# Genetic Identification of Cell Types Underlying Brain Complex Traits Yields Novel Insights Into the Etiology of Parkinson’s Disease

**DOI:** 10.1101/528463

**Authors:** Julien Bryois, Nathan G. Skene, Thomas Folkmann Hansen, Lisette Kogelman, Hunna J. Watson, Zijing Liu, Eating Disorders Working Group of the Psychiatric Genomics Consortium, International Headache Genetics Consortium, 23andMe Research Team, Leo Brueggeman, Gerome Breen, Cynthia M. Bulik, Ernest Arenas, Jens Hjerling-Leffler, Patrick F. Sullivan

## Abstract

Genome-wide association studies (GWAS) have discovered hundreds of loci associated with complex brain disorders, and provide the best current insights into the etiology of these idiopathic traits. However, it remains unclear in which cell types these variants are active, which is essential for understanding etiology and subsequent experimental modeling. Here we integrate GWAS results with single-cell transcriptomic data from the entire mouse nervous system to systematically identify cell types underlying psychiatric disorders, neurological diseases, and brain complex traits. We show that psychiatric disorders are predominantly associated with cortical and hippocampal excitatory neurons, as well as medium spiny neurons from the striatum. Cognitive traits were generally associated with similar cell types but their associations were driven by different genes. Neurological diseases were associated with different cell types, which is consistent with other lines of evidence. Notably, we found that Parkinson’s disease is not only genetically associated with cholinergic and monoaminergic neurons (which include dopaminergic neurons from the substantia nigra) but also with neurons from the enteric system and oligodendrocytes. Using post-mortem brain transcriptomic data, we confirmed alterations in these cells, even at the earliest stages of disease progression. Our study provides an important framework for understanding the cellular basis of complex brain maladies, and reveals an unexpected role of oligodendrocytes in Parkinson’s disease.

## Introduction

Understanding the genetic basis of complex brain disorders is critical for identifying individuals at risk, designing prevention strategies, and developing rational therapeutics. In the last 50 years, twin studies have shown that psychiatric disorders, neurological diseases, and cognitive traits are strongly influenced by genetic factors, explaining a mean of ~50% of the variance in liability ^1^, and GWAS have identified thousands of highly significant loci ^2–5^. However, interpretation of GWAS results remains challenging. First, >90% of the identified variants are located in non-coding regions ^6^, complicating precise identification of risk genes and mechanisms. Second, extensive linkage disequilibrium present in the human genome confounds efforts to pinpoint causal variants and the genes they influence. Finally, it remains unclear in which tissues and cell types these variants are active, and how they disrupt specific biological networks to impact disease risk.

Functional genomic studies from brain are now seen as critical for interpretation of GWAS findings as they can identify functional regions (e.g., open chromatin, enhancers, transcription factor binding sites) and target genes (via chromatin interactions and eQTLs) ^7^. Gene regulation varies substantially across tissues and cell types ^8,9^, and hence it is critical to perform functional genomic studies in empirically identified cell types or tissues.

Multiple groups have developed strategies to identify tissues associated with complex traits ^10–14^, but few have focused on the identification of salient cell types within a tissue. Furthermore, studies aiming to identify relevant cell types often used only a small number of cell types derived from one or few different brain regions ^4,12–18^. For example, we recently showed that, among 24 brain cell types, four types of neurons were consistently associated with schizophrenia ^12^. We were explicit that this conclusion was limited by the relatively few brain regions we studied; other cell types from unsampled regions could conceivably contribute to the disorder.

Here, we integrate a wider range of gene expression data – tissues across the human body and single-cell gene expression data from an entire nervous system – to identify tissues and cell types underlying a large number of complex traits (**Figure 1A,B**). We expand on our prior work by showing that additional cell types are associated with schizophrenia. We also find that psychiatric and cognitive traits are generally associated with similar cell types whereas neurological disorders are associated with different cell types. Notably, we show that Parkinson’s disease is associated with cholinergic and monoaminergic neurons (as expected as these include dopaminergic neurons from the substantia nigra), but also with enteric neurons and oligodendrocytes, providing new clues into its etiology.

**Figure 1:**
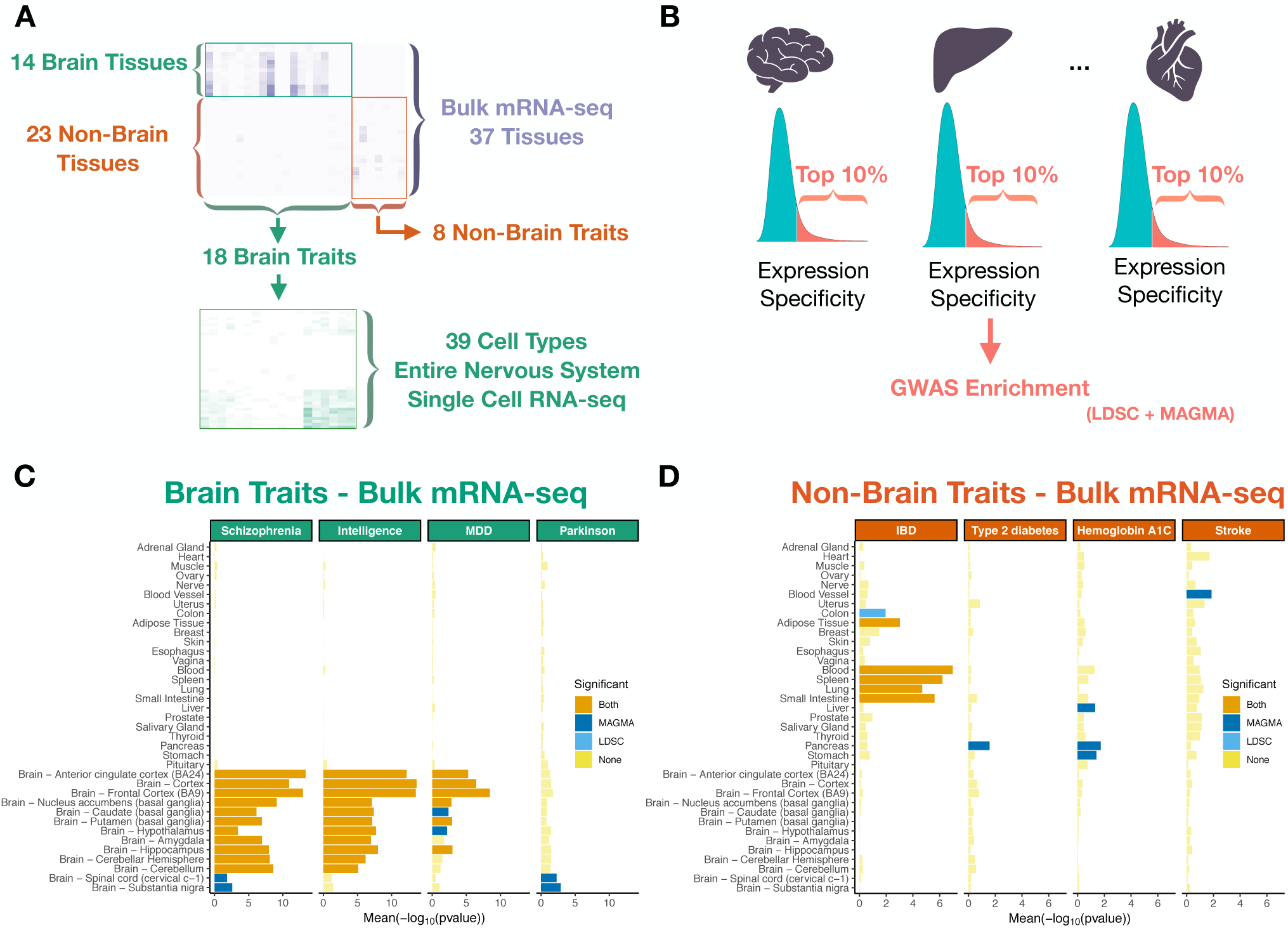
Study design and tissue-level associations. Heat map of trait – tissue/cell types associations (−log_10_P) for the selected traits. (**A**) Trait – tissue/cell types associations were performed using MAGMA and LDSC (testing for enrichment in genetic association of the top 10% most specific genes in each tissue/cell type). (**B**) Tissue – trait associations for selected brain related traits. (**C**) Tissue – trait associations for selected non-brain related traits. (**D**) The mean strength of association (−log_10_P) of MAGMA and LDSC is shown and the bar color indicates whether the tissue is significantly associated with both methods, one method or none (significance threshold: 5% false discovery rate).

## Results

### Genetic correlations among complex traits

Our goal was to use GWAS results to identify relevant tissues and cell types. Our primary focus was human phenotypes whose etiopathology is based in the central nervous system. We thus obtained 18 sets of GWAS summary statistics from European samples for brain-related complex traits. These were selected because they had at least one genome-wide significant association (as of 2018; e.g., Parkinson’s disease, schizophrenia, and IQ). For comparison, we included GWAS summary statistics for 8 diseases and traits with large sample sizes whose etiopathology is not rooted in the central nervous system (e.g., type 2 diabetes). The selection of these conditions allowed contrasts of tissues and cells highlighted by our primary interest in brain phenotypes with non-brain traits. For Parkinson’s disease, we meta-analyzed summary statistics from a published GWAS ^19^ (9,581 cases, 33,245 controls) with self-reported Parkinson’s disease from 23andMe (12,657 cases, 941,588 controls) after finding a high genetic correlation (*r_g_*) ^20^ between the samples (*r_g_*=0.87, s.e=0.068). In this new meta-analysis, we identified 61 independent loci associated with Parkinson’s disease (49 reported previously ^18^ and 12 novel) (**Figure S1**).

We estimated the genetic correlations (*r_g_*) between these 26 traits. We confirmed prior reports ^21,22^ that psychiatric disorders were strongly inter-correlated (e.g., high positive correlations for schizophrenia, bipolar disorder, and MDD) and shared little overlap with neurological disorders (**Figure S2** and **Table S1**). Parkinson’s disease was genetically correlated with intracranial volume ^18^ (*r_g_*=0.29, s.e=0.05) and amyotrophic lateral sclerosis (ALS, *r_g_*=0.19, s.e=0.08), while ALS was negatively correlated with intelligence (*r_g_*=−0.24, s.e=0.06) and hippocampal volume (*r_g_*=−0.24, s.e=0.12). These results indicate that there is substantial genetic heterogeneity across traits, which is a necessary (but not sufficient) condition for trait associations with different tissues or cell types.

### Association of traits with tissues using bulk-tissue RNA-seq

We first aimed to identify the human tissues showing enrichment for genetic associations using bulk-tissue RNA-seq (37 tissues) from GTEx ^8^ (**Figure 1**). To robustly identify the tissues implied by these 26 GWAS, we used two approaches (MAGMA ^23^ and LDSC ^13,24^) which employ different assumptions (**Methods**). For both methods, we tested whether the 10% most specific genes in each tissue were enriched in genetic associations with the different traits (**Figure 1B**).

Examination of non-brain traits found, as expected, associations with salient tissues. For example, as shown in **Figure 1D** and **Table S2**, inflammatory bowel disease was strongly associated with immune tissues (blood, spleen) and alimentary tissues impacted by the disease (small intestine and colon). Lung and adipose tissue were also significantly associated with inflammatory bowel disease, possibly because of the high specificity of immune genes in these two tissues (**Figure S3**). Type 2 diabetes was associated with the pancreas, while hemoglobin A1C, which is used to diagnose type 2 diabetes and monitor glycemic controls in diabetic patients, was associated with the pancreas, liver and stomach (**Figure 1D**). Stroke and coronary artery disease were most associated with blood vessels (**Figure 1D, Figure S4**) and waist to hip ratio was most associated with adipose tissue (**Figure S4**). Thus, our approach can identify the expected tissue associations given the pathophysiology of the different traits.

For brain-related traits (**Figure 1C, S4 and Table S2**), 13 of 18 traits were significantly associated with one or more GTEx brain regions. For example, schizophrenia, intelligence, educational attainment, neuroticism, BMI and MDD were most significantly associated with brain cortex, frontal cortex or anterior cingulate cortex, while Parkinson’s disease was most significantly associated with the substantia nigra (as expected) and spinal cord (**Figure 1C**). Alzheimer’s disease was associated with tissues with prominent roles in immunity (blood and spleen) consistent with other studies ^25–27^, but also with the substantia nigra and spinal cord. Stroke was associated with blood vessel (consistent with a role of arterial pathology in stroke) ^28^. Traits with no or unexpected associations could occur because the primary GWAS had insufficient sample size for its genetic architecture ^29^ or because the tissue RNA-seq data omitted the correct tissue or cell type.

In conclusion, we show that tissue-level gene expression allows identification of relevant tissues for complex traits, indicating that our methodology is suitable to explore trait-gene expression associations at the cell type level.

### Association of brain phenotypes with cell types from the mouse central and peripheral nervous system

We leveraged gene expression data from 39 broad categories of cell types from the mouse central and peripheral nervous system ^30^ to systematically map brain-related traits to cell types (**Figures 2A, S5**). Our use of mouse data to inform human genetic findings was carefully considered (see **Discussion**).

**Figure 2:**
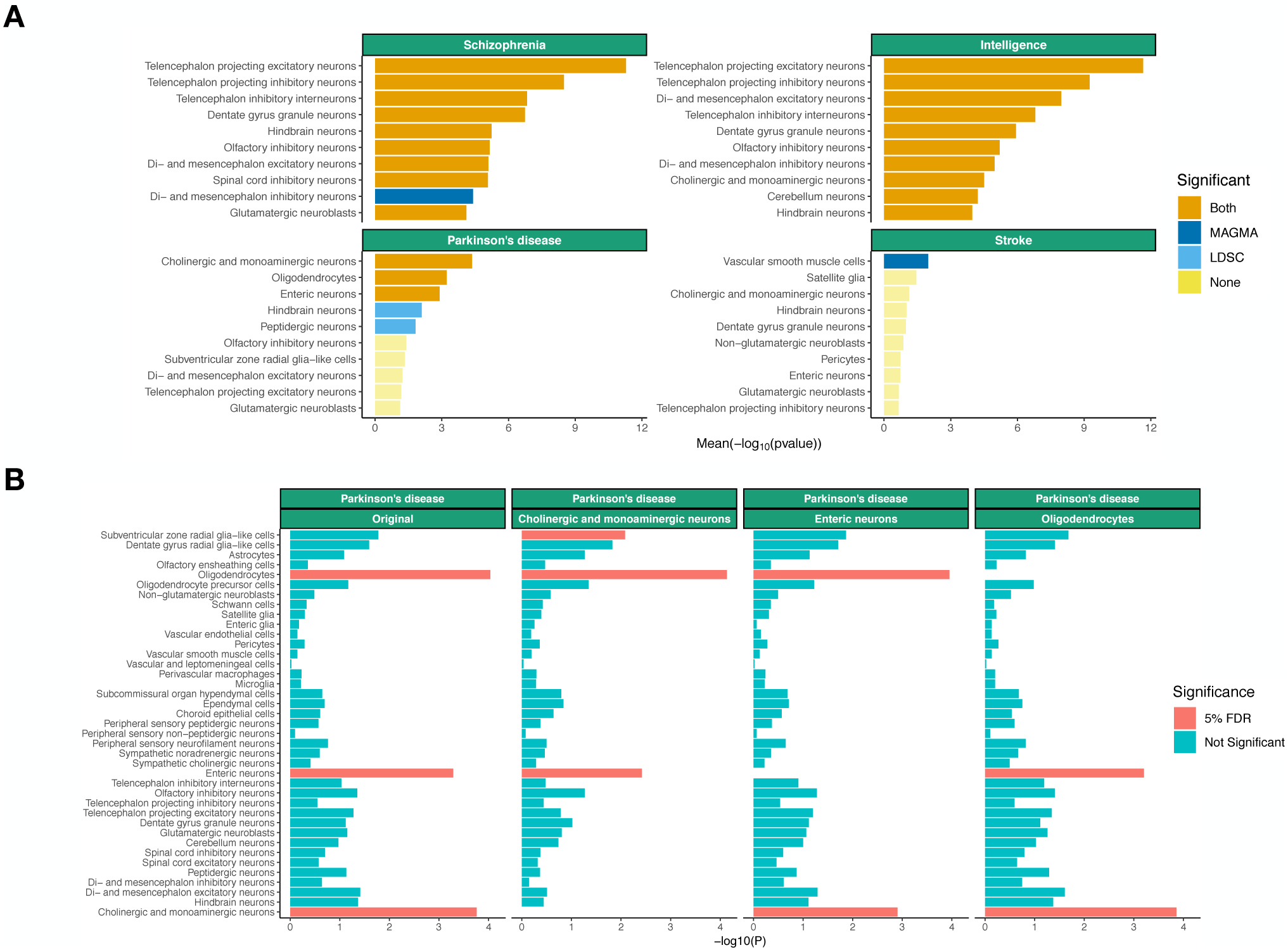
Association of selected brain related traits with cell types from the entire nervous system Associations of the top 10 most associated cell types are shown. (**A**) Conditional analysis results fo Parkinson’s disease using MAGMA. The label indicates the cell type the association analysis is bein conditioned on. (**B**) The mean strength of association (−log_10_P) of MAGMA and LDSC is shown an the bar color indicates whether the cell type is significantly associated with both methods, one metho or none (significance threshold: 5% false discovery rate).

As in our previous study of schizophrenia based on a small number of brain regions ^12^, we found the strongest signals for telencephalon projecting neurons (i.e. excitatory neurons from the cortex, hippocampus and amygdala), telencephalon projecting inhibitory neurons (i.e. medium spiny neurons from the striatum) and telencephalon inhibitory neurons (**Figure 2A** and **Table S3**). We also found that other types of neurons were associated with schizophrenia albeit less significantly (e.g., dentate gyrus granule neurons or hindbrain neurons). Other psychiatric and cognitive traits had similar cellular association patterns to schizophrenia (**Figures S5–6** and **Table S3**). We did not observe any significant associations with immune or vascular cells for any psychiatric disorder or cognitive traits.

Neurological disorders generally implicated fewer cell types, possibly because neurological GWAS had lower signal than GWAS of cognitive, anthropometric, and psychiatric traits (**Figure S7**). Consistent with the genetic correlations reported above, the pattern of associations for neurological disorders was distinct from psychiatric disorders (**Figures S5–6**), again reflecting that neurological disorders have minimal functional overlap with psychiatric disorders ^21^ (**Figure S2**).

Stroke was significantly associated with vascular smooth muscle cells (**Figure 2A**) consistent with an important role of vascular processes for this trait. Amyotrophic lateral sclerosis (a motor neuron disease) was significantly associated with peripheral sensory neurofilament neurons, possibly because of transcriptomic similarities between peripheral sensory and motor neurons (which were not sampled) (**Figure S5**). Alzheimer’s disease had the strongest signal in microglia, as reported previously^11,17,31^, but the association did not survive multiple testing correction.

We found that Parkinson’s disease was significantly associated with cholinergic and monoaminergic neurons (**Figure 2A** and **Table S3**). This cluster consists of neurons (**Table S4**) that are known to degenerate in Parkinson’s disease ^32–34^, such as dopaminergic neurons from the substantia nigra (the hallmark of Parkinson’s disease), but also serotonergic and glutamatergic neurons from the raphe nucleus ^35^, noradrenergic neurons ^36^, as well as neurons from afferent nuclei in the pons ^37^ and the medulla (the brain region associated with the earliest lesions in Parkinson’s disease ^32^). In addition, hindbrain neurons and peptidergic neurons were also significantly associated with Parkinson’s disease (with LDSC only). Therefore, our results capture expected features of Parkinson’s disease and suggest that biological mechanisms intrinsic to these neuronal cell types lead to their selective loss. Interestingly, we also found that enteric neurons were significantly associated with Parkinson’s disease (**Figure 2A**), which is consistent with Braak’s hypothesis, which postulates that Parkinson’s disease could start in the gut and travel to the brain via the vagus nerve ^38,39^. Furthermore, we found that oligodendrocytes (mainly sampled in the midbrain, medulla, pons, spinal cord and thalamus, **Figure S8**) were significantly associated with Parkinson’s disease, indicating a strong glial component to the disorder. This finding was unexpected but consistent with the strong association of the spinal cord at the tissue level (**Figure 1C**), as the spinal cord contains the highest proportion of oligodendrocytes (71%) in the nervous system ^30^. Altogether, these findings provide genetic evidence for a role of enteric neurons, cholinergic and monoaminergic neurons, as well as oligodendrocytes in Parkinson’s disease etiology.

### Neuronal prioritization in the mouse central nervous system

A key goal of this study was to prioritize specific cell types for follow-up experimental studies. As our metric of gene expression specificity was computed based on all cell types in the nervous system, it is possible that the most specific genes in a given cell type capture genes that are shared within a high level category of cell types (e.g. neurons). To rule out this possibility, we computed new specificity metrics based only on neurons from the central nervous system (CNS). We then tested whether the top 10% most specific genes for each CNS neuron were enriched in genetic association for the brain related traits that had a significant association with a CNS neuron (13/18) in our initial analysis.

Using the CNS neuron gene expression specificity metrics, we observed a reduction in the number of neuronal cell types associated with the different traits (**Figure S9**), suggesting that some of the signal was driven by core neuronal genes. For example, the association of telencephalon projecting excitatory neurons with intracranial volume (**Figure S5**) was not significant using the CNS neuron specificity metric (**Figure S9**). However, we found that multiple neuronal cell types remained associated with a number of traits. For example, we found that telencephalon projecting excitatory and projecting inhibitory neurons were strongly associated with schizophrenia, bipolar disorder, educational attainment and intelligence using both LDSC and MAGMA. Similarly, telencephalon projecting excitatory neurons were significantly associated with BMI, neuroticism, MDD, autism and anorexia using one of the two methods (**Figure S9**), while hindbrain neurons and cholinergic and monoaminergic neurons remained significantly associated with Parkinson’s disease (**Figure S9**).

Altogether, these results suggest that specific types of CNS neurons can be prioritized for follow-up experimental studies for multiple traits.

### Cell type-specific and trait-associated genes are enriched in specific biological functions

Understanding which biological functions are dysregulated in different cell types is a key component of the etiology of complex traits. To obtain insights into the biological functions driving cell-type/trait associations, we evaluated GO term enrichment of genes that were specifically expressed (top 20% in a given cell type) and highly associated with a trait (top 10% MAGMA gene-level genetic association). Genes that were highly associated with schizophrenia and specific to telencephalon projecting excitatory neurons were enriched for GO terms related to neurogenesis, synapses, and voltage-gated channels (**Table S5**), suggesting that these functions may be fundamental to schizophrenia. Similarly, genes highly associated with educational attainment, intelligence, bipolar disorder, neuroticism, BMI, anorexia and MDD and highly specific to their most associated cell types were enriched in terms related to neurogenesis, synaptic processes and voltage-gated channels (**Table S5**). In contrast, genes highly associated with stroke and specific to vascular cells were enriched in terms related to vasculature development, while genes highly associated with ALS and peripheral sensory neurofilament neurons were enriched in terms related to lysosomes.

Genes highly associated with Parkinson’s disease and highly specific to cholinergic and monoaminergic neurons were significantly enriched in terms related to endosomes and synapses (**Table S5**). Similarly, genes highly specific to oligodendrocytes and Parkinson’s disease were enriched in endosomes. These results support the hypothesis that the endosomal pathway plays an important role in the etiology of Parkinson’s disease ^40^.

Taken together, we show that cell type-trait associations are driven by genes belonging to specific biological pathways, providing insight into the etiology of complex brain related traits.

### Distinct traits are associated with similar cell types, but through different genes

As noted above, the pattern of associations of psychiatric and cognitive traits were highly correlated across the 39 different cell types tested (**Figure S6**). For example, the Spearman rank correlation of cell type associations (−log_10_P) between schizophrenia and intelligence was 0.96 (0.94 for educational attainment) as both traits had the strongest signal in telencephalon projecting excitatory neurons and little signal in immune or vascular cells. In addition, we observed that genes driving the association signal in the top cell types of the two traits were enriched in relatively similar GO terms involving neurogenesis and synaptic processes. We evaluated two possible explanations for these findings: (a) schizophrenia and intelligence are both associated with the same genes that are specifically expressed in the same cell types or (b) schizophrenia and intelligence are associated with different sets of genes that are both highly specific to the same cell types. Given that these two traits have a significant negative genetic correlation (*r_g_*=−0.22, from GWAS results alone) (**Figure S2** and **Table S1**), we hypothesized that the strong overlap in cell type associations for schizophrenia and intelligence was due to the second explanation.

To evaluate these hypotheses, we tested whether the 10% most specific genes for each cell type were enriched in genetic association for schizophrenia controlling for the gene-level genetic association of intelligence using MAGMA. We found that the pattern of associations were largely unaffected by controlling the schizophrenia cell type association analysis for the gene-level genetic association of intelligence and vice versa (**Figure S10**). Similarly, we found that controlling for educational attainment had little effect on the schizophrenia associations and vice versa (**Figure S11**). In other words, genes driving the cell type associations of schizophrenia appear to be distinct from genes driving the cell types associations of cognitive traits.

### Multiple cell types are independently associated with brain complex traits

Many neuronal cell types passed our stringent significance threshold for multiple brain traits (**Figure 2A** and **S5**). This could be because gene expression profiles are highly correlated across cell types and/or because many cell types are independently associated with the different traits. In order to address this, we performed univariate conditional analysis using MAGMA, testing whether cell type associations remained significant after controlling for the 10% most specific genes from other cell types (**Table S6**). We observed that multiple cell types were independently associated with age at menarche, anorexia, autism, bipolar, BMI, educational attainment, intelligence, MDD, neuroticism and schizophrenia (**Figure S12**). As in our previous study ^12^, we found that the association between schizophrenia and telencephalon projecting inhibitory neurons (i.e. medium spiny neurons) appears to be independent from telencephalon projecting excitatory neurons (i.e. pyramidal neurons). For Parkinson’s disease, we found that enteric neurons, oligodendrocytes and cholinergic and monoaminergic neurons were independently associated with the disorder (**Figure 2B**), suggesting that these three different cell types play an independent role in the etiology of the disorder.

### Replication in other single-cell RNA-seq datasets

To assess the robustness of our results, we repeated these analyses in independent RNA-seq datasets. A key caveat is that these other datasets did not sample the entire nervous system as in the analyses above.

First, we used a single-cell RNA-seq dataset that identified 88 broad categories of cell types (565 subclusters) in 690K single cells from 9 mouse brain regions (frontal cortex, striatum, globus pallidus externus/nucleus basalis, thalamus, hippocampus, posterior cortex, entopeduncular nucleus/subthalamic nucleus, substantia nigra/ventral tegmental area, and cerebellum) ^41^. We found similar patterns of association in this external dataset (**Figure 3A**, **S14** and **Table S7**). Notably, for schizophrenia, we strongly replicated associations with neurons from the cortex, hippocampus and striatum. We also observed similar cell type associations for other psychiatric and cognitive traits (**Figure 3A**, **S13, S14** and **S15**). For neurological disorders, we found that stroke was significantly associated with mural cells while Alzheimer’s disease was significantly associated with microglia (**Figure S14**). The associations of Parkinson’s disease with neurons from the substantia nigra and oligodendrocytes were significant at a nominal level in this dataset (P=0.006 for neurons from the substantia nigra, P=0.027 for oligodendrocytes using LDSC) (**Table S3**). By computing gene expression specificity within neurons, we replicated our previous findings that neurons from the cortex can be prioritized for multiple traits (schizophrenia, bipolar, educational attainment, intelligence, BMI, neuroticism, MDD, anorexia) (**Figure S16**).

**Figure 3:**
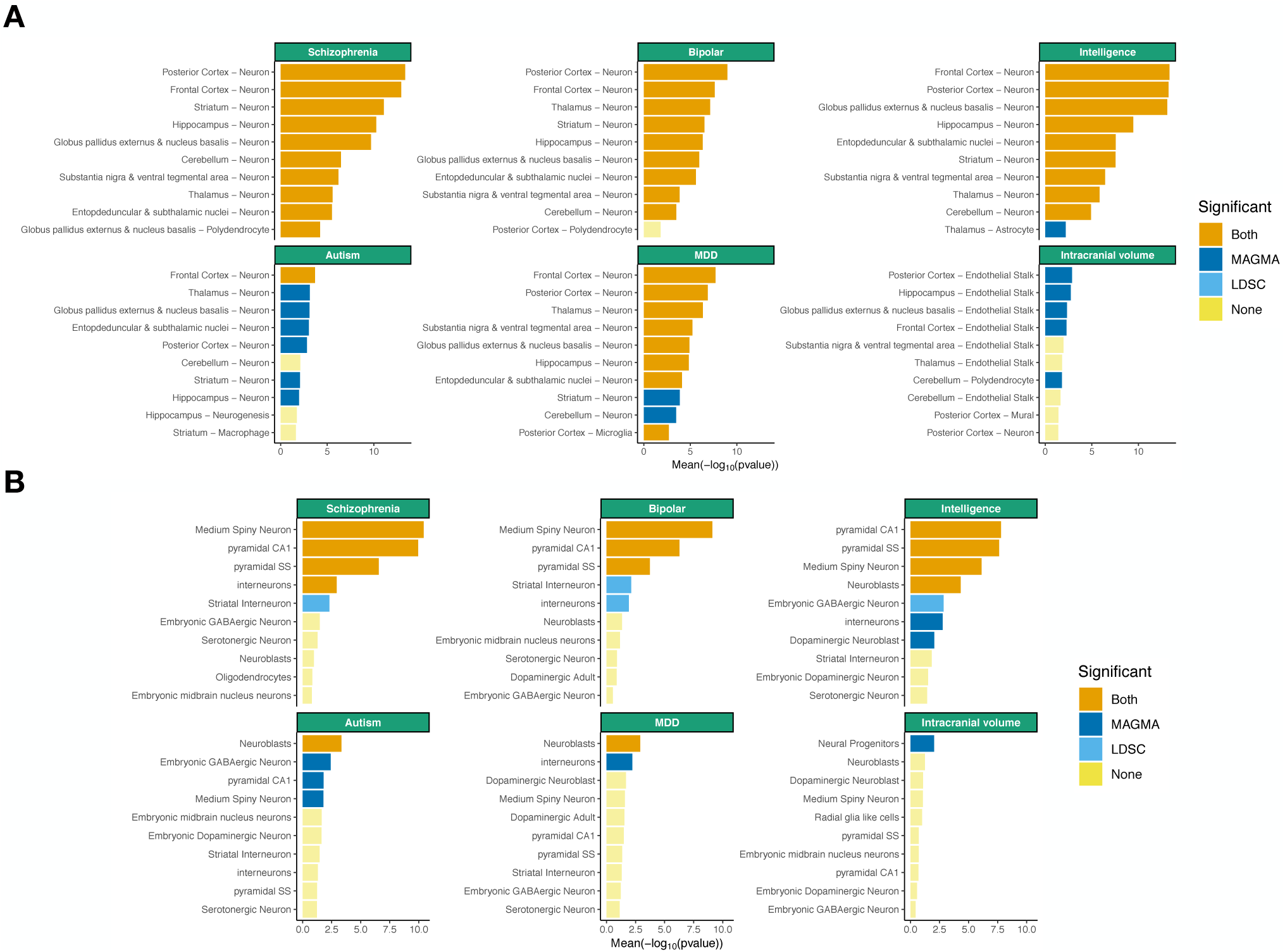
Replication of cell type – trait associations in mouse datasets. Tissue – trait associations are shown for the 10 most association cell types among 88 cell types from 9 different brain regions. (**A**) Tissue – trait associations are shown for the 10 most association cell types among 24 cell types from 5 different brain regions. (**B**) The mean strength of association (−log_10_P) of MAGMA and LDSC is shown and the bar color indicates whether the cell type is significantly associated with both methods, one method or none (significance threshold: 5 % false discovery rate).

Second, we reanalyzed these GWAS datasets using our previous single-cell RNA-seq dataset (24 cell types from the neocortex, hippocampus, striatum, hypothalamus midbrain, and specific enrichments for oligodendrocytes, serotonergic neurons, dopaminergic neurons and cortical parvalbuminergic interneurons, 9970 single cells; **Figure 3B, S17** and **Table S8**). We again found strong associations of pyramidal neurons from the somatosensory cortex, pyramidal neurons from the CA1 region of the hippocampus (both corresponding to telencephalon projecting excitatory neurons in our main dataset), and medium spiny neurons from the striatum (corresponding to telencephalon projecting inhibitory neurons) with psychiatric and cognitive traits. MDD and autism were most associated with neuroblasts, while intracranial volume was most associated with neural progenitors (suggesting that drivers of intracranial volume are cell types implicated in increasing cell mass). The association of dopaminergic adult neurons with Parkinson’s disease was significant at a nominal level using LDSC (P=0.01), while oligodendrocytes did not replicate in this dataset, perhaps because they were not sampled from the regions affected by the disorder (i.e. spinal cord, pons, medulla or midbrain). A within-neuron analysis again found that projecting excitatory (i.e. pyramidal CA1) and projecting inhibitory neurons (i.e. medium spiny neurons) can be prioritized for multiple traits (schizophrenia, bipolar, intelligence, educational attainment, BMI). In addition, we found that neuroblasts could be prioritized for MDD and that neural progenitors could be prioritized for intracranial volume (**Figure S18**) in this dataset.

Third, we evaluated a human single-nuclei RNA-seq dataset consisting of 15 different cell types from cortex and hippocampus ^42^ (**Figure 4A** and **Table S9**). We replicated our findings with psychiatric and cognitive traits being associated with pyramidal neurons (excitatory) and interneurons (inhibitory) from the somatosensory cortex and from the CA1 region of the hippocampus. We also replicated the association of Parkinson’s disease with oligodendrocytes (enteric neurons and cholinergic and monoaminergic neurons were not sampled in this dataset). No cell types reached our significance threshold using specificity metrics computed within-neurons, possibly because of similarities in the transcriptomes of neurons from the cortex and hippocampus.

**Figure 4:**
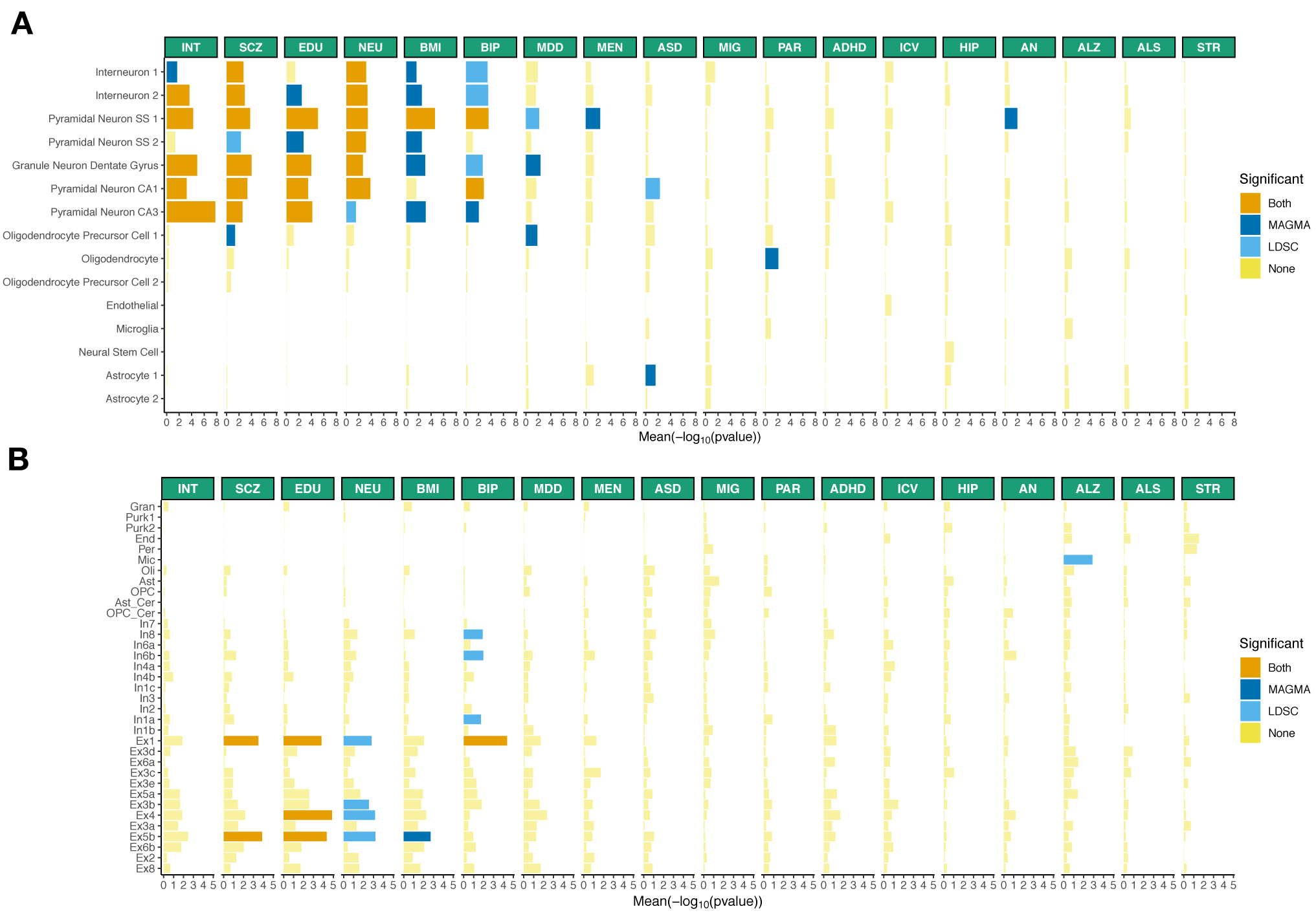
Human replication of cell type – trait associations. Cell type - trait associations for 15 cell types (derived from single-nuclei RNA-seq) from 2 different brain regions (cortex, hippocampus). (**A**) Cell type - trait associations for 31 cell types (derived from single-nuclei RNA-seq) from 3 different brain regions (frontal cortex, visual cortex and cerebellum). (**B**) The mean strength of association (− log_10_P) of MAGMA and LDSC is shown and the bar color indicates whether the cell type is significantly associated with both methods, one method or none (significance threshold: 5% false discovery rate). INT (intelligence), SCZ (schizophrenia), EDU (educational attainment), NEU (neuroticism), BMI (body mass index), BIP (bipolar disorder), MDD (Major depressive disorder), MEN (age at menarche), ASD (autism spectrum disorder), MIG (migraine), PAR (Parkinson’s disease), ADHD (attention deficit hyperactivity disorder), ICV (intracranial volume), HIP (hippocampal volume), AN (anorexia nervosa), ALZ (Alzheimer’s disease), ALS (amyotrophic lateral sclerosis), STR (stroke).

Fourth, we evaluated a human single-nuclei RNA-seq dataset consisting of 31 different cell types from 3 different brain regions (visual cortex, frontal cortex and cerebellum) (**Figure 4B** and **Table S10**). We found that schizophrenia, educational attainment, neuroticism and BMI were associated with excitatory neurons, while bipolar was associated with both excitatory and inhibitory neurons. As observed previously ^11,17,31^, Alzheimer’s disease was significantly associated with microglia. Oligodendrocytes were not significantly associated with Parkinson’s disease in this dataset, again possibly because the spinal cord, pons, medulla and midbrain were not sampled. No cell types reached our significance threshold using specificity metrics computed within neurons in thid dataset.

Most cell type-trait associations were attenuated using human single-nuclei data compared with mouse single-cell RNA-seq data, suggesting that the transcripts that are lost by single-nuclei RNA-seq are important for a large number of disorders and/or that the controlled condition of mouse experiments provide more accurate gene expression quantifications (see **Discussion** and **Figure S19**).

### Comparison with case/control differentially expressed genes at the cell type level

We compared our findings for Alzheimer’s disease (**Table S3**, **Figure 4B, Figure S14**) with a recent study that performed differential expression analysis at the cell type level between 24 Alzheimer’s cases and 24 controls ^43^ (prefrontal cortex, Brodmann area 10). We tested whether the top 500, top 1000 and top 2000 most differentially expressed genes (no pathology vs pathology) in six different cell types (excitatory neurons, inhibitory neurons, oligodendrocytes, oligodendrocytes precursor cells, astrocyte and microglia) were enriched in genetic associations with Alzheimer’s disease using MAGMA. Consistently with our results, we found that genes differentially expressed in microglia were the most associated with Alzheimer’s disease genetics (**Table S11**), indicating that our approach appropriately highlight the relevant cell type at a fraction of the cost of a case-control single cell RNA-seq study. As performing case-control single cell RNA-seq studies in the entire nervous system is currently cost prohibitive, the consistency of our results with the case-control study of Alzheimer’s disease suggests that our results could be leveraged to target specific brain regions and cell types in future case-control genomic studies of brain disorders.

### Validation of oligodendrocyte pathology in Parkinson’s disease

We investigated the role of oligodendrocyte lineage cells in Parkinson’s disease. First, we confirmed the association of oligodendrocytes with Parkinson’s disease by combining evidence across all datasets (Fisher’s combined probability test, P=2.5*10^−7^ using MAGMA and 6.3*10^−3^ using LDSC) (**Table S3** and **Figure S20**). Second, we tested whether oligodendrocytes were significantly associated with Parkinson’s disease conditioning on the top neuronal cell type in the different datasets using MAGMA and found: (a) that oligodendrocytes were independently associated from the top neuronal cell type in our main dataset and in the Habib replication dataset ^42^ at a Bonferroni significant level (P=7.3*10^−5^ and P=1.7*10^−4^ respectively), (b) nominal evidence in the Saunders dataset ^44^ (P=0.018), (c) weak evidence in the Skene ^12^ (P=0.12) and Lake ^45^ datasets (P=0.2) and (d) combining the conditional evidence from all datasets, oligodendrocytes were significantly associated with Parkinson’s disease independently of the top neuronal association (P=1.2*10^−7^, Fisher’s combined probability test).

Third, we tested whether genes with rare variants associated with Parkinsonism (**Table S12**) were specifically expressed in cell types from the mouse nervous system (**Method**). As for the common variant, we found the strongest enrichment for cholinergic and monoaminergic neurons (**Table S13**). However, we did not observe any significant enrichments for oligodendrocytes or enteric neurons using genes associated with rare variants in Parkinsonism.

Fourth, we applied EWCE ^11^ to test whether genes that are up/down-regulated in human post-mortem Parkinson’s disease brains (from six separate cohorts) were enriched in cell types located in the substantia nigra and ventral midbrain (**Figure 5**). Three of the studies had a case-control design and measured gene expression in: (a) the substantia nigra of 9 controls and 16 cases ^46^, (b) the medial substantia nigra of 8 controls and 15 cases ^47^, and (c) the lateral substantia nigra of 7 controls and 9 cases ^47^. In all three studies, downregulated genes in Parkinson’s disease were specifically enriched in dopaminergic neurons (consistent with the loss of this particular cell type in disease), while upregulated genes were significantly enriched in cells from the oligodendrocyte lineage. This suggests that an increased oligodendrocyte activity or proliferation could play a role in Parkinson’s disease etiology. Surprisingly, no enrichment was observed for microglia, despite recent findings ^48,49^.

**Figure 5:**
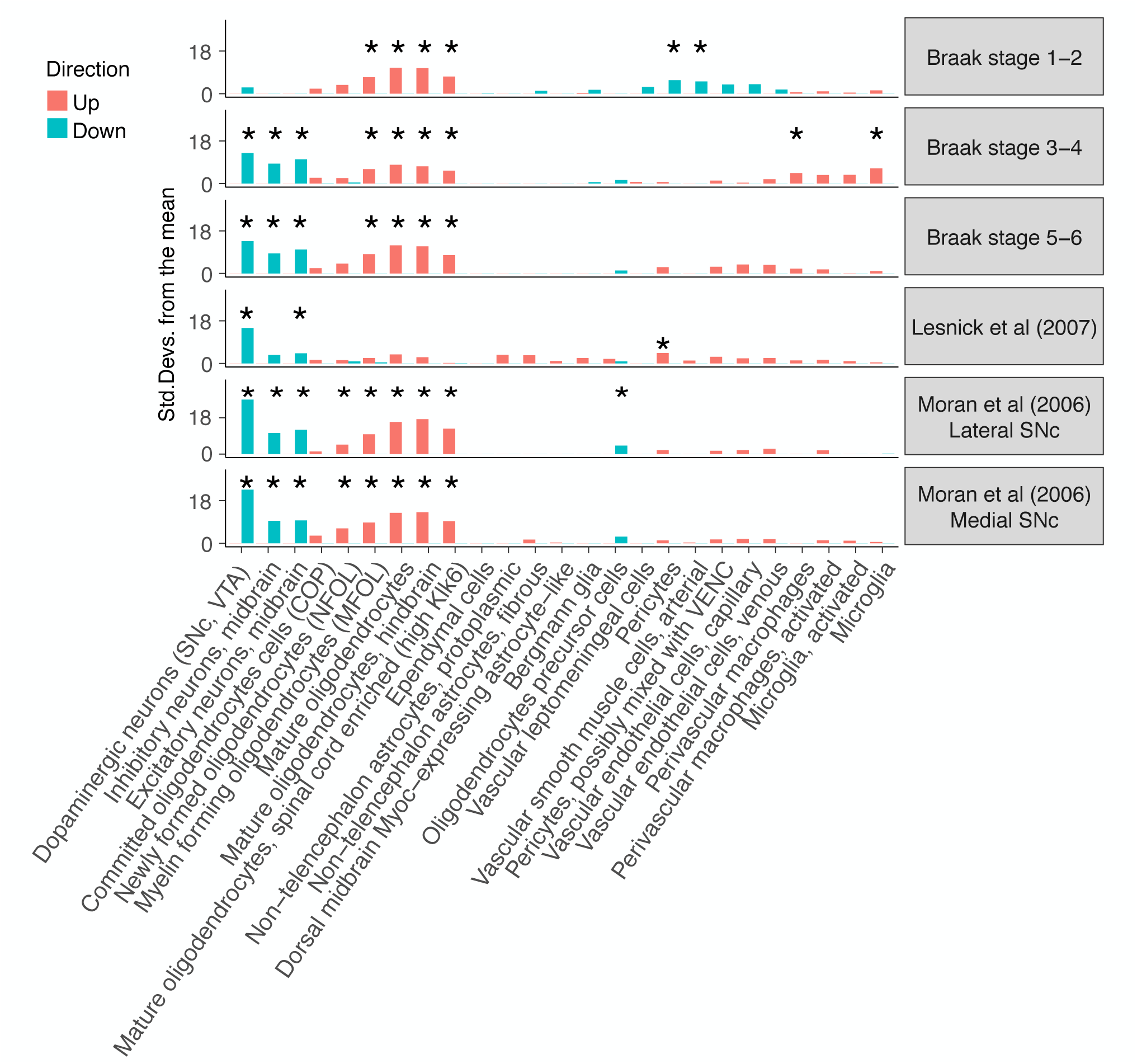
Enrichment of Parkinson’s disease differentially expressed genes in cell types from the substantia nigra. Enrichment of the 500 most up/down regulated genes (Braak stage 0 vs Braak stage 1—2, 3—4 and 5—6, as well as cases vs controls) in postmortem human substantia nigra gene expression samples. The enrichments were obtained using EWCE^11^. A star shows significant enrichments after multiple testing correction (P<0.05/(25*6).

We also analyzed gene expression data from post-mortem human brains which had been scored by neuropathologists for their Braak stage ^50^. Differential expression was calculated between brains with Braak scores of zero (controls) and brains with Braak scores of 1—2, 3—4 and 5—6. At the latter stages (Braak scores 3—4 and 5—6), downregulated genes were specifically expressed in dopaminergic neurons, while upregulated genes were specifically expressed in oligodendrocytes (**Figure 5**), as observed in the case-control studies. Moreover, Braak stage 1 and 2 are characterized by little degeneration in the substantia nigra and, consistently, we found that downregulated genes were not enriched in dopaminergic neurons at this stage. Notably, upregulated genes were already strongly enriched in oligodendrocytes at Braak Stages 1-2. These results not only support the genetic evidence indicating that oligodendrocytes may play a causal role in Parkinson’s disease, but indicate that their involvement precedes the emergence of pathological changes in the substantia nigra.

## Discussion

In this study, we used gene expression data from cells sampled from the entire nervous system to systematically map cell types to GWAS results from multiple psychiatric, cognitive, and neurological complex phenotypes.

We note several limitations. First, we again emphasize that we can implicate a particular cell type but it is premature to exclude cell types for which we do not have data ^12^. Second, we used gene expression data from mouse to understand human phenotypes. We believe our approach is appropriate for several reasons. (A) Crucially, the key findings replicated in human data. (B) Single-cell RNA-seq is achievable in mouse but difficult in human neurons (where single-nuclei RNA-seq is typical ^42,45,51,52^). In brain, differences between single-cell and single-nuclei RNA-seq are important as transcripts that are missed by sequencing nuclei are important for psychiatric disorders, and we previously showed that dendritically-transported transcripts (important for schizophrenia) are specifically depleted from nuclei datasets ^12^ (we confirmed this finding in four additional datasets, **Figure S19**). (C) Correlations in gene expression for cell type across species is high (median correlation 0.68, **Figure S21**), and as high or higher than correlations across methods within cell type and species (single-cell vs single-nuclei RNA-seq, median correlation 0.6) ^53^. (D) We evaluated protein-coding genes with 1:1 orthologs between mouse and human. These constitute most human protein-coding genes, and these genes are generally highly conserved particularly in the nervous system. We did not study genes present in one species but not in the other. (E) More specifically, we previously showed that gene expression data cluster by cell type and not by species ^12^, indicating broad conservation of core brain cellular functions across species. (F) We used a large number of genes to map cell types to traits (~1500 genes for each cell type), minimizing potential bias due to individual genes differentially expressed across species. (G) If there were strong differences in cell type gene expression between mouse and human, we would not expect that specific genes in mouse cell types would be enriched in genetic associations with human disorders. However, it remains possible that some cell types have different gene expression patterns between mouse and human, are only present in one species, have a different function or are involved in different brain circuits.

A third limitation is that gene expression data were from adolescent mice. Although many psychiatric and neurological disorders have onsets in adolescence, some have onsets earlier (autism) or later (Alzheimer’s and Parkinson’s disease). It is thus possible that some cell types are vulnerable at specific developmental times. Data from studies mapping cell types across brain development and aging are required to resolve this issue.

For schizophrenia, we replicated and extended our previous findings ^12^. We found the most significant associations for neurons located in the cortex, hippocampus and striatum (**Figure 2A, 3**) in multiple independent datasets, and showed that these neuronal cell types can be prioritized among neurons (**Figure S9, S16** and **S18**). These results are consistent with the strong schizophrenia heritability enrichment observed in open chromatin regions from: human dorsolateral prefrontal cortex ^54^; human cortical, striatal and hippocampal neurons ^55^; and mouse open chromatin regions from cortical excitatory and inhibitory neurons ^56^. This degree of replication in independent transcriptomic datasets from multiple groups along with consistent findings using orthogonal open chromatin data is notable, and strongly implicates these cell types in the etiology of schizophrenia.

Moreover, we found that other psychiatric traits implicated largely similar cell types. These biological findings are consistent with genetic and epidemiological evidence of a general psychopathy factor underlying diverse clinical psychiatric disorders ^21,57,58^. Although intelligence and educational attainment implicated similar cell types, conditional analyses showed that the same cell types were implicated for different reasons. This suggests that different sets of genes highly specific to the same cell types contribute independently to schizophrenia and cognitive traits.

A number of studies have argued that the immune system plays a causal role in some psychiatric disorders ^59,60^. Our results did not implicate any brain immune cell types in psychiatric disorders. We interpret these negative findings cautiously as we did not fully sample the immune system. It is also possible that a small number of genes are active in immune cell types and that these cell types play an important role in the etiology of psychiatric disorders. Finally, if immune functions are salient for a small subset of patients, GWAS may not identify these loci without larger and more detailed studies.

Our findings for neurological disorders were strikingly different from psychiatric disorders. In contrast to previous studies that either did not identify any cell type associations with Parkinson’s disease ^61^ or identified significant associations with cell types from the adaptive immune system ^49^, we found that cholinergic and monoaminergic neurons (which include dopaminergic neurons), enteric neurons and oligodendrocytes were significantly and independently associated with the disease. It is well established that loss of dopaminergic neurons in the substantia nigra is a hallmark of Parkinson’s disease. Our findings suggest that dopaminergic neuron loss in Parkinson’s disease is at least partly due to intrinsic biological mechanisms. In addition, other type of cholinergic and monoaminergic neurons are known to degenerate in Parkinson’s disease (e.g., raphe nucleus serotonergic neurons and cholinergic neurons of the pons), suggesting that specific pathological mechanisms may be shared across these neurons and lead to their degeneration. Two theories for the selective vulnerability of neuronal populations in Parkinson’s disease currently exist: the “spread Lewy pathology model” which assumes cell-to-cell contacts enabling spreading of prion-like α-synuclein aggregates ^62^; and the “threshold theory” ^63,64^ which proposes that the vulnerable cell types degenerate due to molecular/functional biological similarities in a cell-autonomous fashion. While both theories are compatible and can co-exist, our findings support the existence of cell autonomous mechanisms contributing to selective vulnerability. We caution that we do not know if all cholinergic and monoaminergic neurons show degeneration or functional impairment. However, analysis of the cellular mechanisms driving the association of cholinergic and monoaminergic neurons with Parkinson’s disease revealed endosomal trafficking as a plausible common pathogenic mechanism.

Interestingly, enteric neurons were also associated with Parkinson’s disease. This result is in line with prior evidence implicating the gut in Parkinson’s disease. Notably, dopaminergic defects and Lewy bodies (i.e. abnormal aggregates of proteins enriched in α-synuclein) are found in the enteric nervous system of patients affected by Parkinson’s disease ^65,66^. In addition, Lewy bodies have been observed in patients up to 20 years prior to their diagnosis ^67^ and sectioning of the vagus nerve (which connects the enteric nervous system to the central nervous system) was shown to reduce the risk of developing Parkinson’s disease ^68^. Therefore, our results linking enteric neurons with Parkinson’s disease provides new genetic evidence for Braak’s hypothesis, which postulates that Parkinson’s disease could start in the gut, travel along the vagus nerve, and affect the brain years after disease initiation^38^.

The association of oligodendrocytes with Parkinson’s disease was more unexpected. A possible explanation is that this association could be due to a related disorder (e.g., multiple system atrophy, characterized by Parkinsonism and accumulation of α-synuclein in glial cytoplasmic inclusions ^69^). However, this explanation is unlikely as multiple system atrophy is a very rare disorder; hence, only a few patients are likely to have been included in the Parkinson’s disease GWAS which could not have affected the GWAS results. In addition, misdiagnosis is unlikely to have led to the association of Parkinson’s disease with oligodendrocytes. Indeed, we found a high genetic correlation between self-reported diagnosis from the 23andMe cohort and a previous GWAS of clinically-ascertained Parkinson’s disease ^19^. In addition, self-report of Parkinson’s disease in 23andMe subjects was confirmed by a neurologist in all 50 cases evaluated ^70^.

We did not find an association of oligodendrocytes with Parkinsonism for genes affected by rare variants. This result may reflect etiological differences between sporadic and familial forms of the disease or the low power and insufficient number of genes tested. Prior evidence has suggested an involvement of oligodendrocytes in Parkinson’s disease. For example, α-synuclein-containing inclusions have been reported in oligodendrocytes in Parkinson’s disease brains ^71^. These inclusions (“coiled bodies”) are typically found throughout the brainstem nuclei and fiber tracts ^72^. Although the presence of coiled bodies in oligodendrocytes is a common, specific, and well-documented neuropathological feature of Parkinson’s disease, the importance of this cell type and its early involvement in disease has not been fully recognized. Our findings suggest that intrinsic genetic alterations in oligodendrocytes occur at an early stage of disease, which precedes the emergence of neurodegeneration in the substantia nigra, arguing for a key role of this cell type in Parkinson’s disease etiology.

Taken together, we integrated genetics and single-cell gene expression data from the entire nervous system to systematically identify cell types underlying brain complex traits. We believe that this a critical step in the understanding of the etiology of brain disorders and that these results will guide modelling of brain disorders and functional genomic studies.

## Methods

### GWAS results

Our goal was to use GWAS results to identify relevant tissues and cell types. Our primary focus was human phenotypes whose etiopathology is based in the central nervous system. We thus obtained 18 sets of GWAS summary statistics from European samples for brain-related complex traits. These were selected because they had at least one genome-wide significant association (as of 2018; e.g., Parkinson’s disease, schizophrenia, and IQ). For comparison, we included GWAS summary statistics for 8 diseases and traits with large sample sizes whose etiopathology is not rooted in the central nervous system (e.g., type 2 diabetes). The selection of these conditions allowed contrasts of tissues and cells highlighted by our primary interest in brain phenotypes with non-brain traits.

The phenotypes were: schizophrenia ^2^, educational attainment ^3^, intelligence ^15^, body mass index ^5^, bipolar disorder ^73^, neuroticism ^4^, major depressive disorder ^74^, age at menarche ^75^, autism ^76^, migraine^77^, amyotrophic lateral sclerosis ^78^, ADHD ^79^, Alzheimer’s disease ^26^, age at menopause ^80^, coronary artery disease ^81^, height ^5^, hemoglobin A1c ^82^, hippocampal volume ^83^, inflammatory bowel disease^84^, intracranial volume ^85^, stroke ^86^, type 2 diabetes mellitus ^87^, type 2 diabetes adjusted for BMI ^87^, waist-hip ratio adjusted for BMI ^88^, and anorexia nervosa ^89^.

For Parkinson’s disease, we performed an inverse variance-weighted meta-analysis ^90^ using summary statistics from Nalls et al. ^19^ (9,581 cases, 33,245 controls) and summary statistics from 23andMe (12,657 cases, 941,588 controls). We found a very high genetic correlation (*r_g_*) ^20^ between results from these cohorts (*r_g_*=0.87, s.e=0.068) with little evidence of sample overlap (LDSC bivariate intercept=0.0288, s.e=0.0066). The P-values from the meta-analysis strongly deviated from the expected (**Figure S22**) but was consistent with polygenicity (LDSC intercept=1.0048, s.e=0.008) rather than uncontrolled inflation ^20^.

### Gene expression data

We collected publicly available single-cell RNA-seq data from different studies. The core dataset of our analysis is a study that sampled more than 500K single cells from the entire mouse nervous system (19 regions) and identified 39 broad categories (level 4) and 265 refined cell types (level 5)^30^. The 39 cell types expressed a median of 16417 genes, had a median UMI total count of ~8.6M and summed the expression of a median of 1501 single cells (**Table S14**). The replication datasets were: 1) a mouse study that sampled 690K single cells from 9 brain regions and identified 565 cell types ^91^ (note that we averaged the UMI counts by broad categories of cell type in each brain region, resulting in 88 different cell types); 2) our prior mouse study of ~10K cells from 5 different brain regions (and samples enriched for oligodendrocytes, dopaminergic neurons, serotonergic neurons and cortical parvalbuminergic interneurons) that identified 24 broad categories and 149 refined cell types ^12^; 3) a study that sampled 19,550 nuclei from frozen adult human post-mortem hippocampus and prefrontal cortex and identified 16 cell types ^42^; 4) a study that generated 36,166 single-nuclei expression measurements (after quality control) from the human visual cortex, frontal cortex and cerebellum ^45^. We also obtained bulk tissues RNA-seq gene expression data from 53 tissues from the GTEx consortium ^8^ (v8, median across samples).

### Gene expression data processing

All datasets were processed uniformly. First we computed the mean expression for each gene in each cell type from the single-cell expression data (if this statistics was not provided by the authors). We used the pre-computed median expression across individuals for the GTEx dataset and excluded tissues that were not sampled in at least 100 individuals, non-natural tissues (e.g. EBV-transformed lymphocytes) and testis (outlier using hierarchical clustering). We then averaged the expression of tissues by organ (with the exception of brain tissues) resulting in gene expression profiles of a total of 37 tissues. For all datasets, we filtered out any genes with non-unique names, genes not expressed in any cell types, non-protein coding genes, and, for mouse datasets, genes that had no expert curated 1:1 orthologs between mouse and human (Mouse Genome Informatics, The Jackson laboratory, version 11/22/2016). Gene expression was then scaled to a total of 1M UMIs (or transcript per million (TPM)) for each cell type/tissue. We then calculated a metric of gene expression specificity by dividing the expression of each gene in each cell type by the total expression of that gene in all cell types, leading to values ranging from 0 to 1 for each gene (0: meaning that the gene is not expressed in that cell type, 0.6: that 60% of the total expression of that gene is performed in that cell type, 1: that 100% of the expression of that gene is performed in that cell type). The top 10% most specific genes (**Table S15** and **Table S16**) in each tissue/cell type partially overlapped for related tissues/cell types, did not overlap for unrelated tissue/cell types and allowed to cluster related tissues/cell types as expected (**Figure S23** and **Figure S24**).

### MAGMA primary and conditional analyses

MAGMA (v1.06b) ^23^ is a software for gene-set enrichment analysis using GWAS summary statistics. Briefly, MAGMA computes a gene-level association statistic by averaging P-values of SNPs located around a gene (taking into account LD structure). The gene-level association statistic is then transformed to a Z-value. MAGMA can then be used to test whether a gene set is a predictor of the gene-level association statistic of the trait (Z-value) in a linear regression framework. MAGMA accounts for a number of important covariates such as gene size, gene density, mean sample size for tested SNPs per gene, the inverse of the minor allele counts per gene and the log of these metrics.

For each GWAS summary statistics, we excluded any SNPs with INFO score <0.6, with MAF < 1% or with estimated odds ratio > 25 or smaller than 1/25, the MHC region (chr6:25-34 Mb) for all GWAS and the *APOE* region (chr19:45020859–45844508) for the Alzheimer’s GWAS. We set a window of 35kb upstream to 10kb downstream of the gene coordinates to compute gene-level association statistics and used the European reference panel from the phase 3 of the 1000 genomes project ^92^ as the reference population. For each trait, we then used MAGMA to test whether the 10% most specific gene in each tissue/cell type was associated with gene-level genetic association with the trait. Only genes with at least 1TPM or 1 UMI per million in the tested cell type were used for this analysis. The significance level of the different cell types was highly correlated with the effect size of the cell type (**Figure S25**) with values ranging between 0.999 and 1 across the 18 brain related traits in the Zeisel et al. dataset ^93^. The significance threshold was set to a 5% false discovery rate across all tissues/cell types and traits within each dataset.

MAGMA can also perform conditional analyses given its linear regression framework. We used MAGMA to test whether cell types were associated with a specific trait conditioning on the gene-level genetic association of another trait (Z-value from MAGMA .out file) or to look for associations of cell types conditioning on the 10% most specific genes from other cell types by adding these variables as covariate in the model.

To test whether MAGMA was well-calibrated, we randomly permuted the gene labels of the schizophrenia gene-level association statistic file a thousand times. We then looked for association between the 10% most specific genes in each cell type and the randomized gene-level schizophrenia association statistics. We observed that MAGMA was slightly conservative with less than 5% of the random samplings having a P-value <0.05 (**Figure S26**).

We also evaluated the effect of varying window sizes (for the SNPs to gene assignment step of MAGMA) on the schizophrenia cell type associations strength (−log_10_(P)). We observed strong Pearson correlations in cell type associations strength (−log_10_(P)) across the different window sizes tested (**Figure S27**). Our selected window size (35kb upstream to 10 kb downstream) had Pearson correlations ranging from 0.94 to 0.98 with the other window sizes, indicating that our results are robust to this parameter.

In a recent paper, Watanabe et al. ^94^ introduced a different methodology to test for cell type – complex trait association based on MAGMA. Their proposed methodology tests for a positive relationship between gene expression levels and gene-level genetic associations with a complex trait (using all genes). Their method uses the average expression of each gene in all cell types in the dataset as a covariate. We examined the method of Watanabe et al. in detail, and decided against its use for multiple reasons.

First, Watanabe et al. hypothesize that genes with higher levels of expression should be more associated with a trait. In extended discussions among our team (which include multiple neuroscientists), we have strong reservations about the appropriateness and biological meaningfulness of this hypothesis; it is a strong requirement and is at odds with decades of neuroscience research where molecules expressed a low levels can have profound biological impact. For instance, many cell-type specific genes that are disease relevant are expressed at moderate levels (e.g., *Drd2* is in the 10% most specific genes in telencephalon projecting inhibitory neurons but in the bottom 30% of expression levels). Our method does not make this hypothesis.

Second, the method of Watanabe et al. corrects for the average expression of all cell types in a dataset. This practice is, in our view, problematic as it necessarily forces dependence on the composition of a scRNA-seq dataset. For instance, if a dataset consists mostly of neurons, this amounts to correcting for neuronal expression and necessarily erodes power to detect trait enrichment in neurons. Alternatively, if a dataset is composed mostly of non-neuronal cells, this will impacts the detection of enrichment in non-neuronal cells.

Third, preliminary results indicate that the method of Watanabe et al. is sensitive to scaling. As different cell types express different numbers of genes, scaling to the same total read counts affects the average gene expression across cell types (which they use as a covariate), leading to different results with different choices of scaling factors (e.g., scaling to 10k vs 1 million reads). Our method is not liable to this issue.

### LD score regression analysis

We used partitioned LD score regression ^95^ to test whether the top 10% most specific genes of each cell type (based on our specificity metric described above) were enriched in heritability for the diverse traits. Only genes with at least 1TPM or 1 UMI per million in the tested cell type were used for this analysis. In order to capture most regulatory elements that could contribute to the effect of the region on the trait, we extended the gene coordinates by 100kb upstream and by 100kb downstream of each gene as previously ^13^. SNPs located in 100kb regions surrounding the top 10% most specific genes in each cell type were added to the baseline model (consisting of 53 different annotations) independently for each cell type (one file for each cell type). We then selected the coefficient z-score p-value as a measure of the association of the cell type with the traits. The significance threshold was set to a 5% false discovery rate across all tissues/cell types and traits within each dataset. All plots show the mean −log_10_(P) of partitioned LDscore regression and MAGMA. All results for MAGMA or LDSC are available in supplementary data files.

We evaluated the effect of varying window sizes and varying the percentage of most specific genes on the schizophrenia cell type associations strength (−log_10_P). We observed strong Pearson correlations in cell type associations strength (−log_10_P) across the different percentage and window sizes tested (**Figure S28**). Our selected window size (100 kb upstream to 100 kb downstream, top 10% most specific genes) had Pearson correlations ranging from 0.96 to 1 with the other window sizes and percentage, indicating that our results are robust to these parameters.

### MAGMA vs LDSC ranking

In order to test whether the cell type ranking obtained using MAGMA and LDSC in the Zeisel et al. dataset ^30^ were similar, we computed the Spearman rank correlation of the cell types association strength (−log_10_P) between the two methods for each complex trait. The Spearman rank correlation was strongly correlated with *λ_GC_* (a measure of the deviation of the GWAS test statistics from the expected) (Spearman ρ=0.89) (**Figure S29**) and with the average number of cell types below our stringent significance threshold (Spearman ρ=0.92), indicating that the overall ranking of the cell types is very similar between the two methods, provided that the GWAS is well powered (**Figure S30**). In addition, we found that *λ_GC_* was strongly correlated with the strength of association of the top tissue (−log_10_P) (Spearman ρ=0.88) (**Figure S31**), as well as with the effect size (beta) of the top tissue (Spearman ρ=0.9), indicating that cell type – trait associations are stronger for well powered GWAS. The significance level (−log_10_P) was also strongly correlated with the effect size (Spearman ρ=0.996) (**Figure S31**) for the top cell type of each trait.

### Dendritic depletion analysis

This analysis was performed as previously described ^12^. In brief, all datasets were reduced to a set of six common cell types: pyramidal neurons, interneurons, astrocytes, microglia and oligodendrocyte precursors. Specificity was recalculated using only these six cell types. Comparisons were then made between pairs of datasets (denoted in the graph with the format ‘X versus Y’). The difference in specificity for a set of dendrite enriched genes is calculated between the datasets. Differences in specificity are also calculated for random sets of genes selected from the background gene set. The probability and z-score for the difference in specificity for the dendritic genes is thus estimated. Dendritically enriched transcripts were obtained from Supplementary Table 10 of Cajigas et al. ^96^. For the KI dataset ^12^, we used S1 pyramidal neurons. For the Zeisel 2018 dataset ^30^ we used all ACTE* cells as astrocytes, TEGLU* as pyramidal neurons, TEINH* as interneurons, OPC as oligodendrocyte precursors and MGL* as microglia. For the Saunders dataset ^41^, we used all Neuron.Slc17a7 cellt ypes from FC, HC or PC as pyramidal neurons; all Neuron.Gad1Gad2 cell types from FC, HC or PC as interneurons; Polydendrocye as OPCs; Astrocyte as astrocytes, and Microglia as microglia. The Lake datasets both came from a single publication ^45^ which had data from frontal cortex, visual cortex and cerebellum. The cerebellum data was not used here. Data from frontal and visual cortices were analyzed separately. All other datasets were used as described in our previous publication ^12^. The code and data for this analysis are available as an R package (see code availability below).

### GO term enrichment

We tested whether genes that were highly specific to a trait-associated cell type (top 20% in a given cell type) AND highly associated with the genetics of the traits (top 10% MAGMA gene-level genetic association) were enriched in biological functions using the *topGO* R package ^97^. As background, we used genes that were highly specific to the cell type (top 20%) OR highly associated with the trait (top 10% MAGMA gene-level genetic association).

### Parkinson’s disease rare variant enrichments

We searched the literature for genes associated with Parkinsonism on the basis of rare and familial mutations. We found 66 genes (listed in **Table S12**). We used linear regression to test whether the z-scaled specificity metric (per cell type) of the 66 genes were greater than 0 in the different cell types.

### Parkinson’s disease post-mortem transcriptomes

The Moran dataset ^47^ was obtained from GEO (accession GSE8397). Processing of the U133a and U133b Cel files was done separately. The data was read in using the ReadAffy function from the R affy package ^98^, then Robust Multi-array Averaging (RMA) was applied. The U133a and U133b array expression data were merged after applying RMA. Probe annotations and mapping to HGNC symbols was done using the biomaRt R package ^99^. Differential expression analysis was performed using limma ^100^ taking age and gender as covariates. The Lesnick dataset ^46^ was obtained from GEO (accession GSE7621). Data was processed as for the Moran dataset: however, age was not available to use as a covariate. The Disjkstra dataset ^50^ was obtained from GEO (accession GSE49036) and processed as above: the gender and RIN values were used as covariates. As the transcriptome datasets measured gene expression in the substantia nigra, we only kept cell types that are present in the substantia nigra or ventral midbrain for our EWCE ^11^ analysis. We computed a new specificity matrix based on the substantia nigra or ventral midbrain cells from the Zeisel dataset (level 5) using EWCE ^11^. The EWCE analysis was performed on the 500 most up or down regulated genes using 10,000 bootstrapping replicates.

## Code availability

The code used to generate these results is available at: https://github.com/jbryois/scRNA_disease. An R package for performing cell type enrichments using magma is also available from: https://github.com/NathanSkene/MAGMA_Celltyping.

## Data availability

All single-cell expression data are publicly available. Most summary statistics used in this study are publicly available. The migraine GWAS can be obtained by contacting the authors ^77^. The Parkinson’s disease summary statistics from 23andMe can be obtained under an agreement that protects the privacy of 23andMe research participants (https://research.23andme.com/collaborate/#publication).

## Acknowledgments

J.B. was funded by a grant from the Swiss National Science Foundation (P400PB_180792). N.G.S. was supported by the Wellcome Trust (108726/Z/15/Z). N.G.S and L.B. performed part of the work at the Systems Genetics of Neurodegeneration summer school funded by BMBF as part of the e:Med program (FKZ 01ZX1704). J.H.-L. was funded by the Swedish Research Council (Vetenskapsrådet, award 2014-3863), StratNeuro, the Wellcome Trust (108726/Z/15/Z) and the Swedish Brain Foundation (Hjärnfonden). PFS was supported by the Swedish Research Council (Vetenskapsrådet, award D0886501), the Horizon 2020 Program of the European Union (COSYN, RIA grant agreement n° 610307), and US NIMH (U01 MH109528 and R01 MH077139). KH was supported by The Michael J. Fox Foundation for Parkinson’s Research (grant MJFF12737). EA was supported by the Swedish Research Council (VR 2016-01526), Swedish Foundation for Strategic Research (SLA SB16-0065), Karolinska Institutet (SFO Strat. Regen., Senior grant 2018), Cancerfonden (CAN 2016/572), Hjärnfonden (FO2017-0059) and Chen Zuckeberg Initiative: Neurodegeneration Challenge Network (2018-191929-5022). CMB acknowledges funding from the Swedish Research Council (Vetenskapsrådet, award: 538-2013-8864) and the Klarman Family Foundation. We thank the research participants from 23andMe and other cohorts for their contribution to this study.

Members of the 23andMe Research Team: Michelle Agee, Babak Alipanahi, Adam Auton, Robert K. Bell, Katarzyna Bryc, Sarah L. Elson, Pierre Fontanillas, Nicholas A. Furlotte, Karl Heilbron, David A. Hinds, Karen E. Huber, Aaron Kleinman, Nadia K. Litterman, Jennifer C. McCreight, Matthew H. McIntyre, Joanna L. Mountain, Elizabeth S. Noblin, Carrie A.M. Northover, Steven J. Pitts, J. Fah Sathirapongsasuti, Olga V. Sazonova, Janie F. Shelton, Suyash Shringarpure, Chao Tian, Joyce Y. Tung, Vladimir Vacic, and Catherine H. Wilson.

## Author contributions

J.B., N.G.S., J.H.-L. and P.F.S. designed the study, wrote and reviewed the manuscript; J.B performed the analyses pertaining to Figure 1–4, Figure S1–S18, Figure S20–S31, table S1-S11 and table S13-S16; N.G.S performed the analyses pertaining to Figure 5, Figure S19 and table S12-S13; T.F.H, L.K. and the I.H.G.C provided the migraine GWAS summary statistics; H.W., the E.D.W.G.P.G.C, G.B. and C.M.B performed the anorexia GWAS; Z.L. contributed to the revision of the manuscript, The 23andMe R.T. provided GWAS summary statistics for Parkinson’s disease in the 23andMe cohort. L.B. contributed to the post-mortem differential expression analysis (Figure 5); E.A. and K.H. provided expert knowledge on Parkinson’s disease and reviewed the manuscript.

## Tables

**Table S1:** Genetic correlations across traits

**Table S2:** Association P-value between GTEx tissues and all traits

**Table S3:** Association P-value between cell types from the entire mouse nervous system and all traits (Zeisel et al. 2018)

**Table S4:** Sub clusters of cell types corresponding to the 39 broad categories of cell types across the mouse nervous system

**Table S5:** GO term enrichment of genes highly specific to cell type and diseases

**Table S6:** Univariate conditional analysis results using MAGMA

**Table S7:** Association P-value between cell types from 9 mouse brain regions and all traits (Saunders et al. 2018)

**Table S8:** Association P-value between cell types from 5 mouse brain regions and all traits (Skene et al. 2018)

**Table S9:** Association P-value between cell types from 2 human brain regions and all traits (Habib et al. 2017)

**Table S10:** Association P-value between cell types from 3 human brain regions and all traits (Lake et al. 2018)

**Table S11:** Association of Alzheimer’s disease differentially expressed genes in 6 different cell types with Alzheimer’s common variant genetics using MAGMA.

**Table S12:** Rare and familial genetic mutations associated with Parkinsonism

**Table S13:** Cell type enrichment results using rare and familial genetic mutations associated with Parkinsonism. The one-sided pvalues were computed using linear regression, testing whether the average specificity metric of the gene set was higher than 0 (z-scaled specificity metrics per tissue).

**Table S14:** Summary statistics of cell types from the mouse nervous system (Zeisel et al. 2018)

**Table S15:** Top 10% most specific genes per tissue for the GTEx dataset

**Table S16:** Top 10% most specific genes per cell type for the Zeisel dataset

## Supplementary Figures

**Figure S1:**
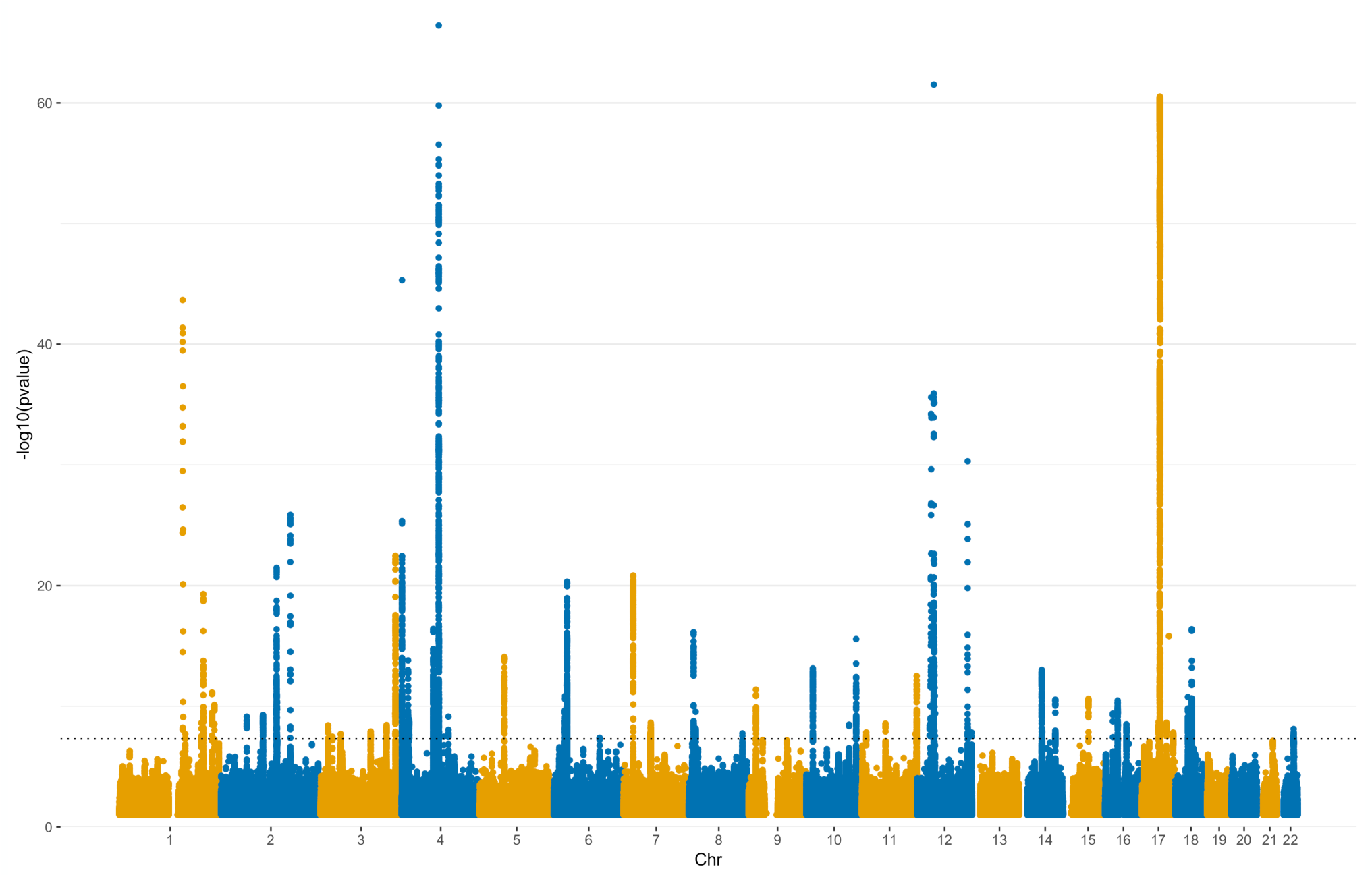
Manhattan plot of Parkinson’s disease meta-analysis. The black dotted line represents the genome-wide significance threshold (5×10^−8^).

**Figure S2:**
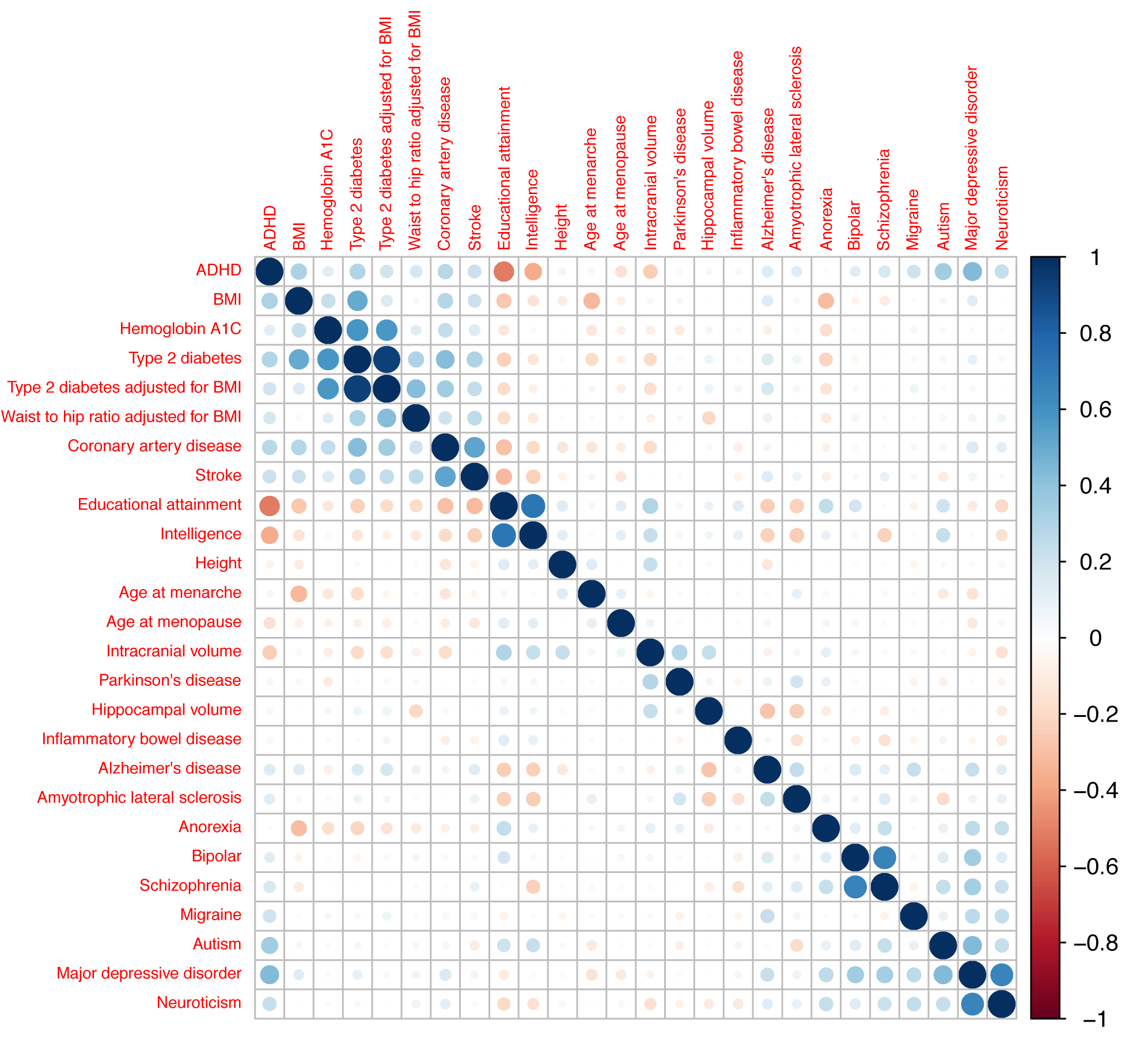
Genetic correlation across traits. The genetic correlation across traits were computed using LDSC^101^. Traits are ordered based on hierarchical clustering.

**Figure S3:**
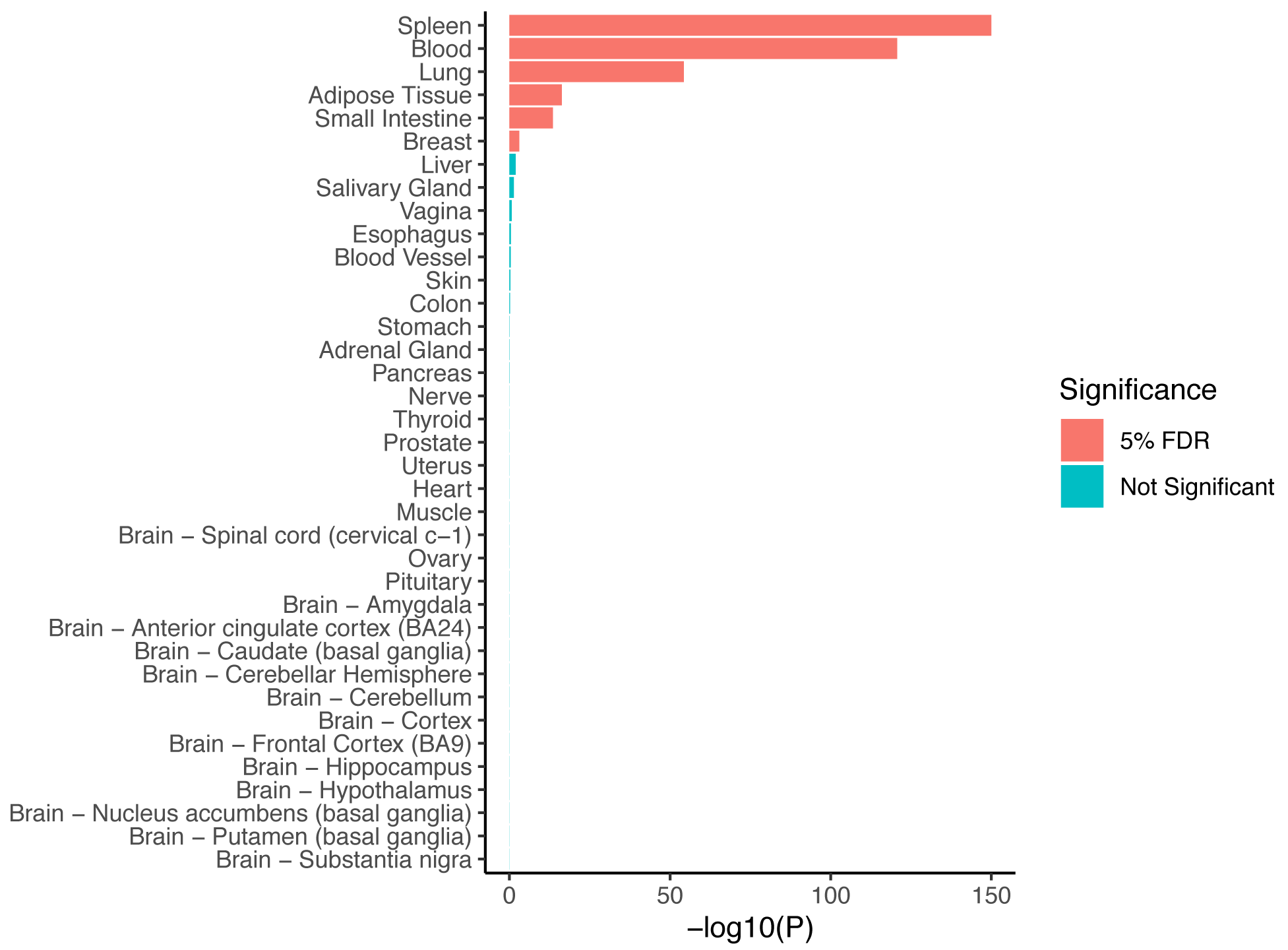
Enrichment of immune genes in GTEx tissues. Enrichment pvalues of genes belonging to the GO term “Immune System Process” in the 10% most specific genes in each tissue. The one-sided pvalues were computed using linear regression, testing whether the average specificity metric of the gene set was higher than 0 (z-scaled specificity metrics per tissue). The GO term was selected because it is the most associated with inflammatory bowel disease using MAGMA.

**Figure S4:**
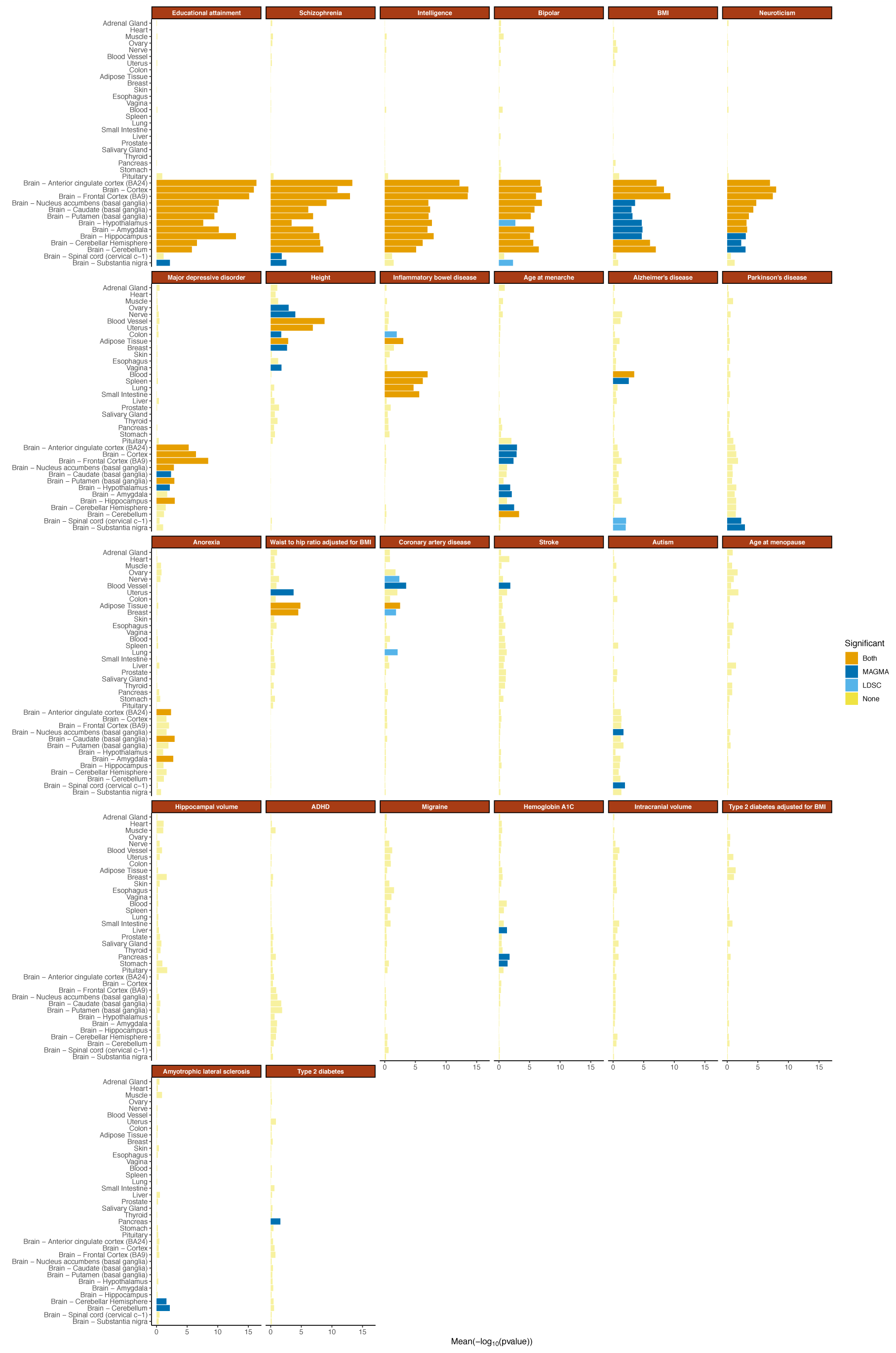
Tissue – trait associations for all traits. The mean strength of association (−log_10_P) of MAGMA and LDSC is shown and the bar color indicates whether the tissue is significantly associated with both methods, one method or none (significance threshold: 5% false discovery rate).

**Figure S5:**
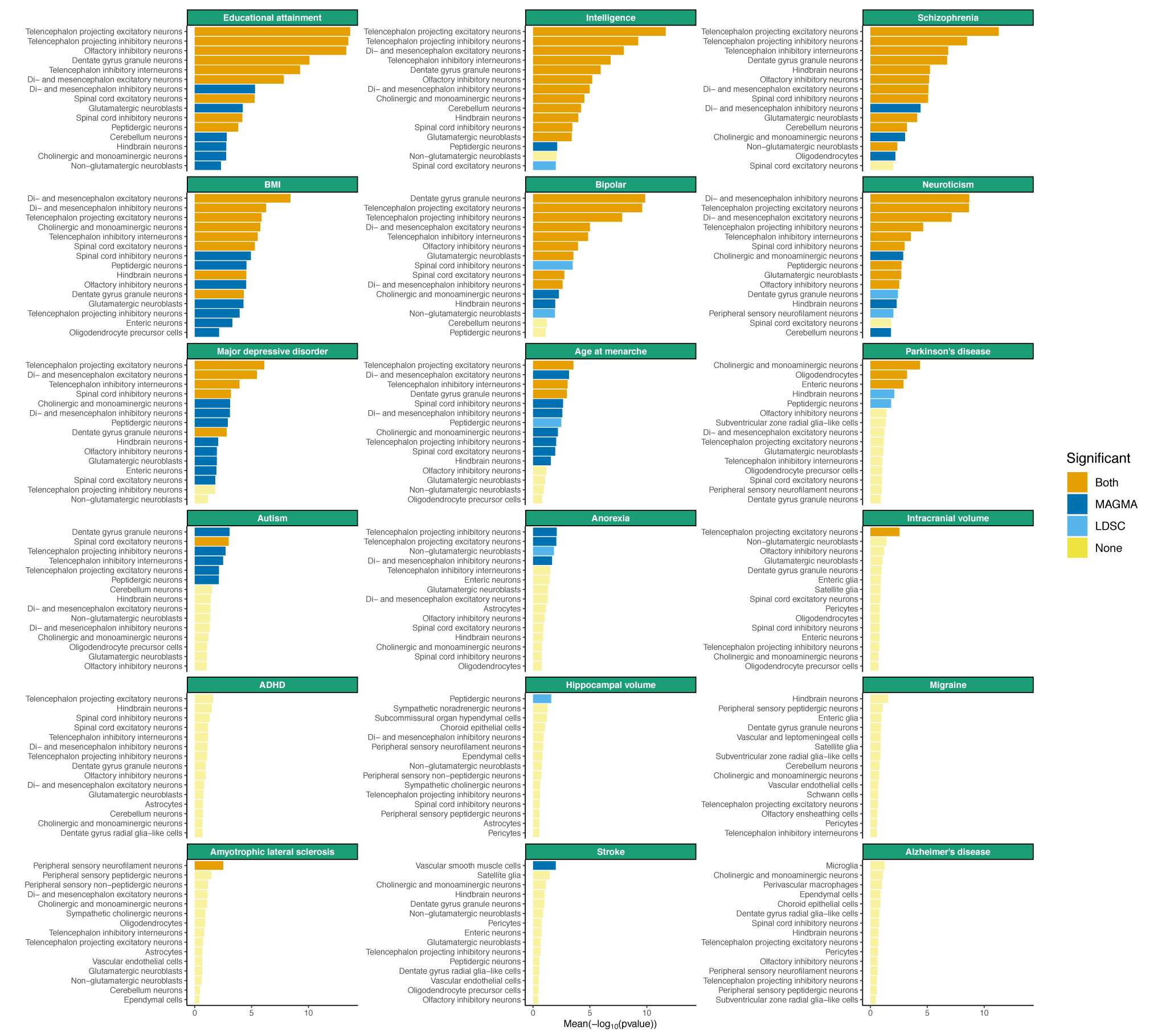
Associations of brain related traits with cell types from the entire mouse nervous system. Associations of the top 15 most associated cell types are shown. The mean strength of association (− log_10_P) of MAGMA and LDSC is shown and the bar color indicates whether the cell type is significantly associated with both methods, one method or none (significance threshold: 5% false discovery rate).

**Figure S6:**
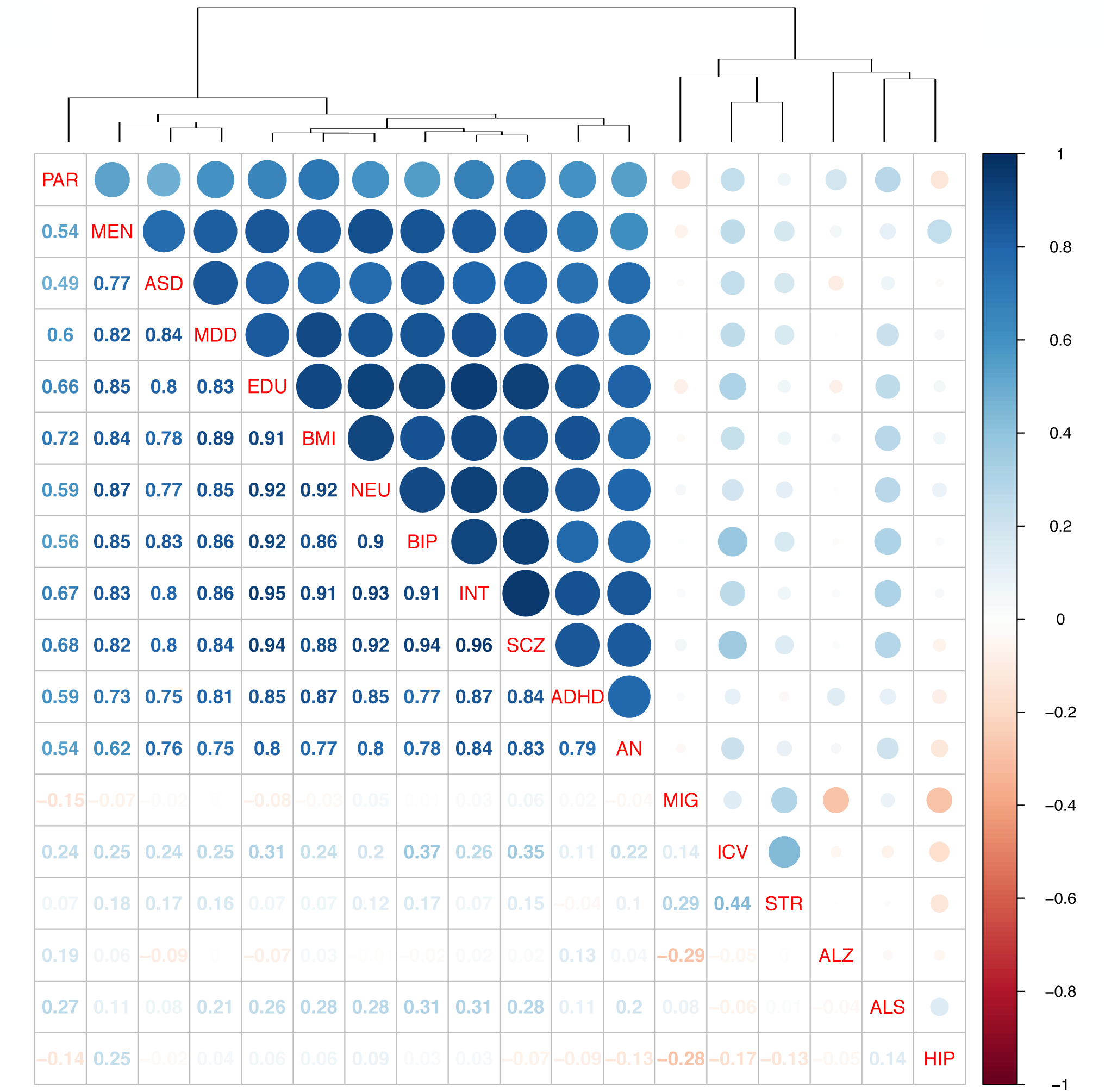
Correlation in cell type associations across traits. The Spearman rank correlations between the cell types associations across traits (−log_10_P) are shown. SCZ (schizophrenia), EDU (educational attainment), INT (intelligence), BMI (body mass index), BIP (bipolar disorder), NEU (neuroticism), PAR (Parkinson’s disease), MDD (Major depressive disorder), MEN (age at menarche), ICV (intracranial volume), ASD (autism spectrum disorder), STR (stroke), AN (anorexia nervosa), MIG (migraine), ALS (amyotrophic lateral sclerosis), ADHD (attention deficit hyperactivity disorder), ALZ (Alzheimer’s disease), HIP (hippocampal volume).

**Figure S7:**
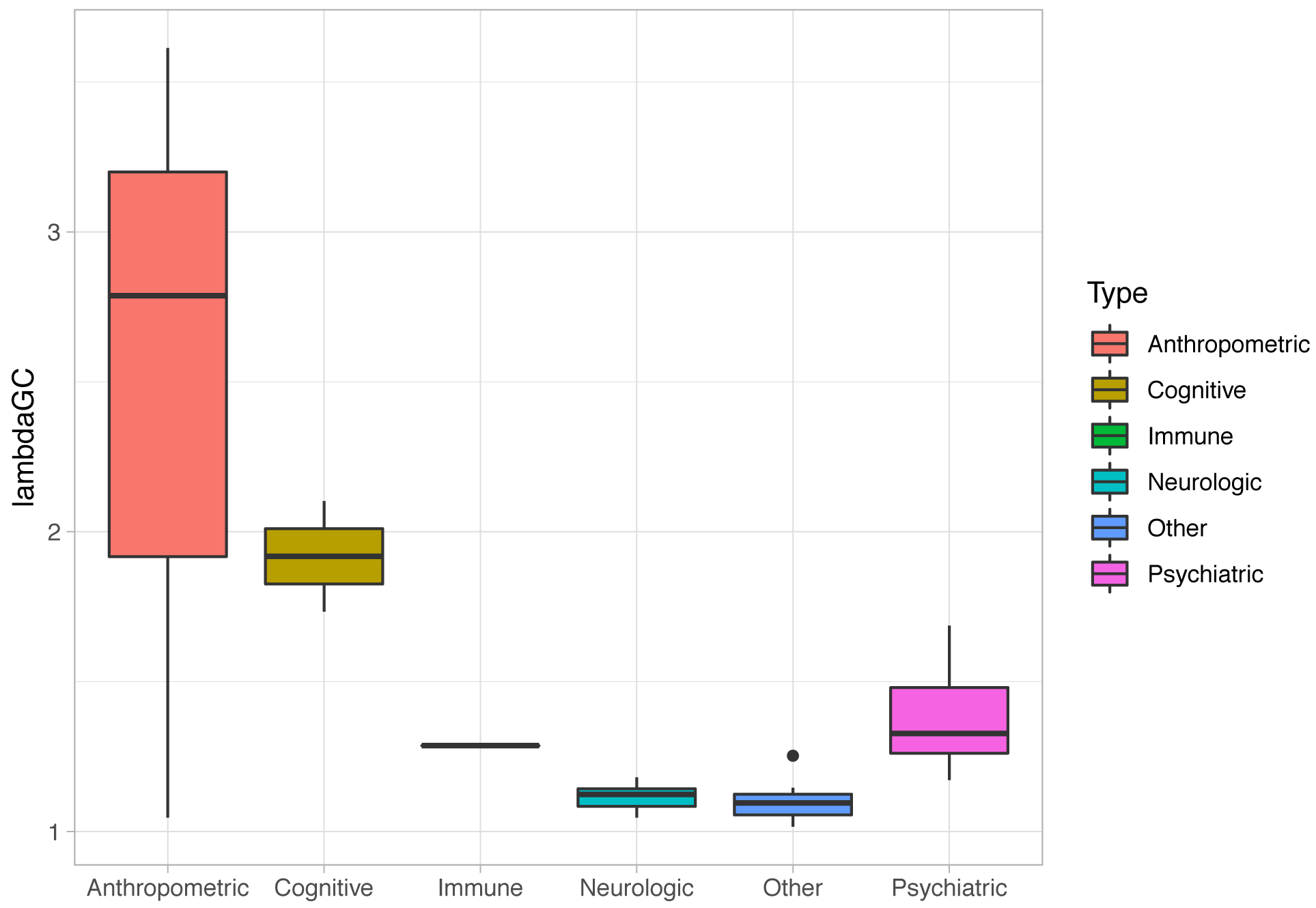
GWAS signal to noise ratio (λ_GC_) by category of GWAS trait. Boxplot of the λ _GC_ of the different GWAS by category of trait. λ_GC_ was estimated using LDSC for each GWAS.

**Figure S8:**
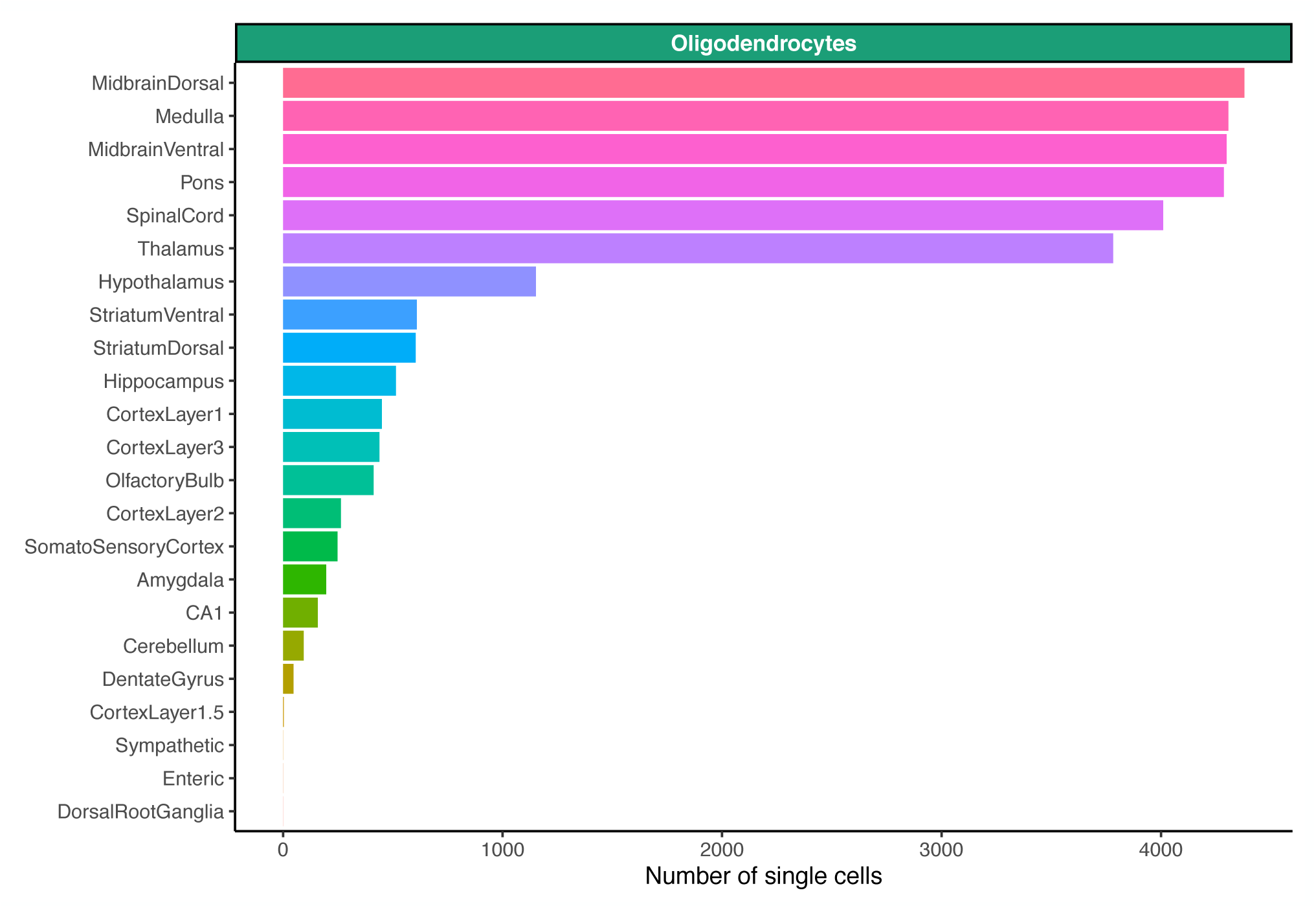
Number of single cells forming the oligodendrocyte cluster. Number of single cells per region of the mouse nervous system used to estimate the average gene expression of oligodendrocytes.

**Figure S9:**
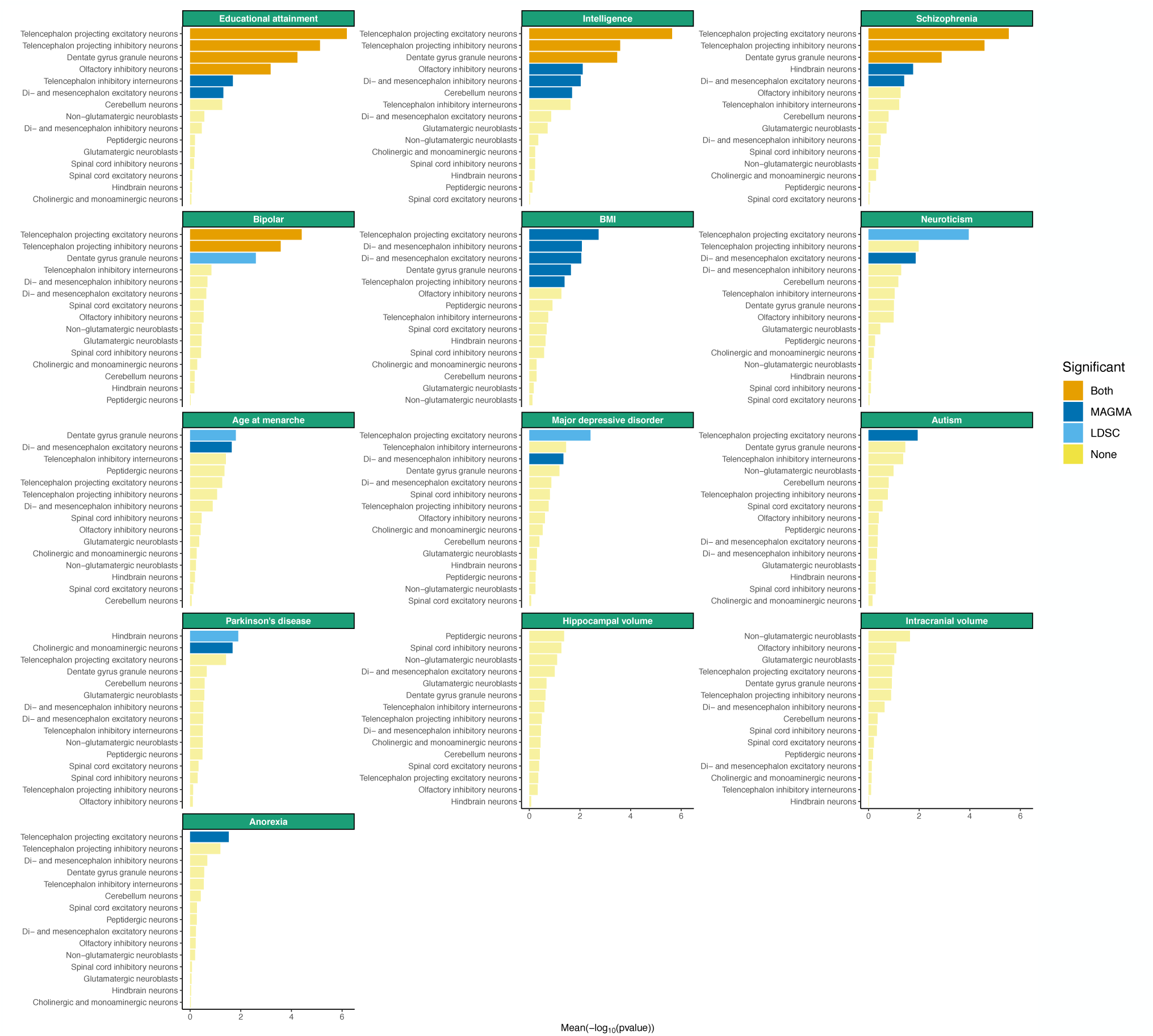
Associations of brain related traits with neurons from the central nervous system. Associations of the 15 most associated neurons from the central nervous system (CNS) are shown. The specificity metrics were computed only using neurons from the CNS. The mean strength of association (−log_10_P) of MAGMA and LDSC is shown and the bar color indicates whether the cell type is significantly associated with both methods, one method or none (significance threshold: 5% false discovery rate).

**Figure S10:**
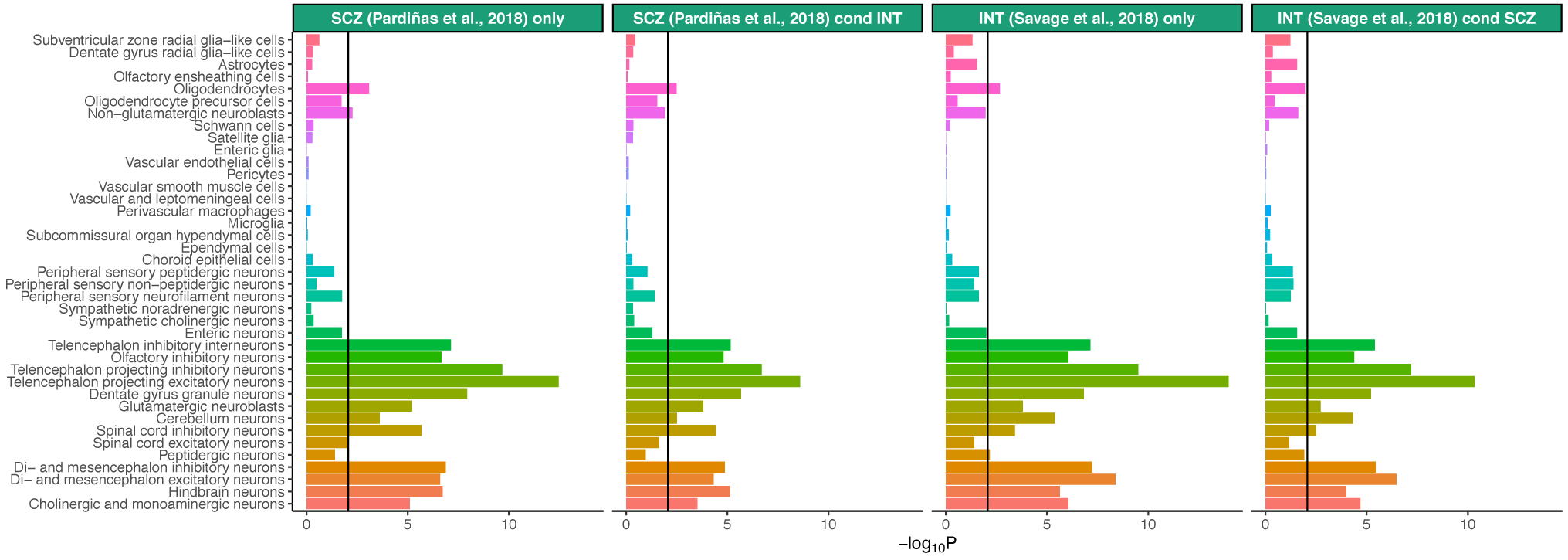
Associations of cell types with schizophrenia/intelligence conditioning on gene-level genetic association of intelligence/schizophrenia. MAGMA association strength for each cell type before and after conditioning on gene-level genetic association for another trait. The black bar represents the significance threshold (5% false discovery rate). SCZ (schizophrenia), INT (intelligence).

**Figure S11:**
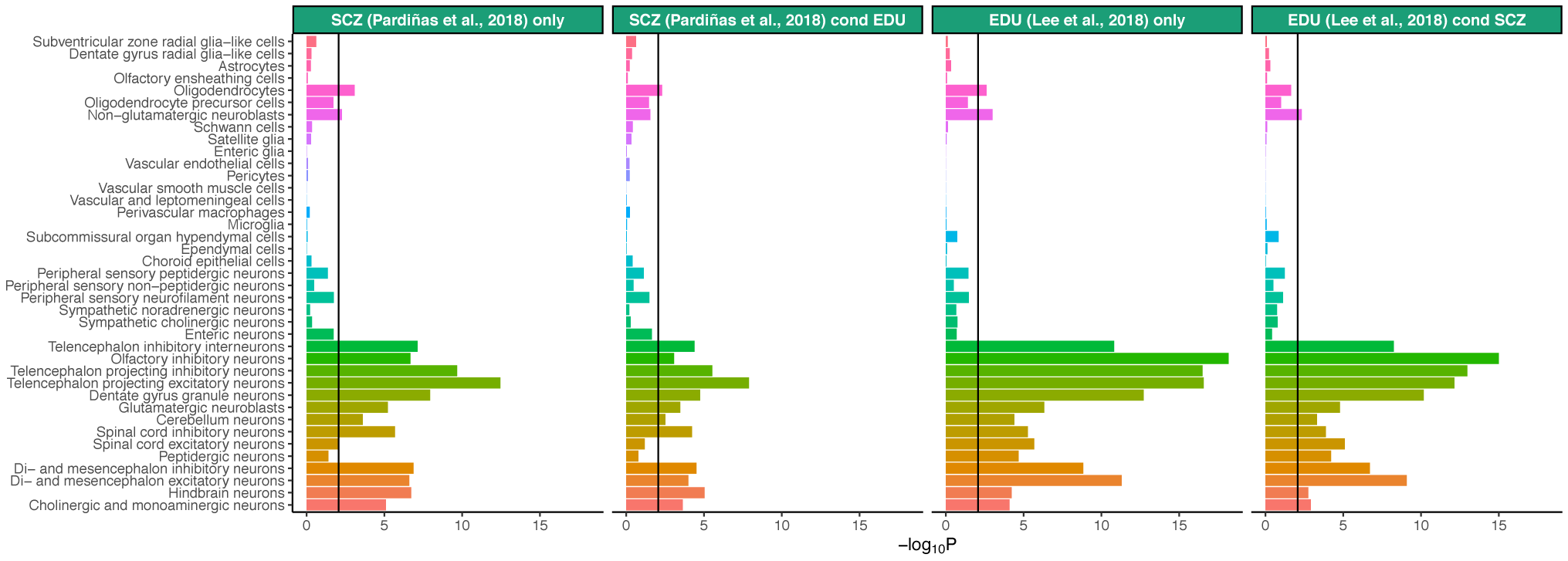
Associations of cell types with schizophrenia/educational attainment conditioning on gene-level genetic association of educational attainment/schizophrenia. MAGMA association strength for each cell type before and after conditioning on gene-level genetic association for another trait. The black bar represents the significance threshold (5% false discovery rate). SCZ (schizophrenia), EDU (educational attainment).

**Figure S12:**
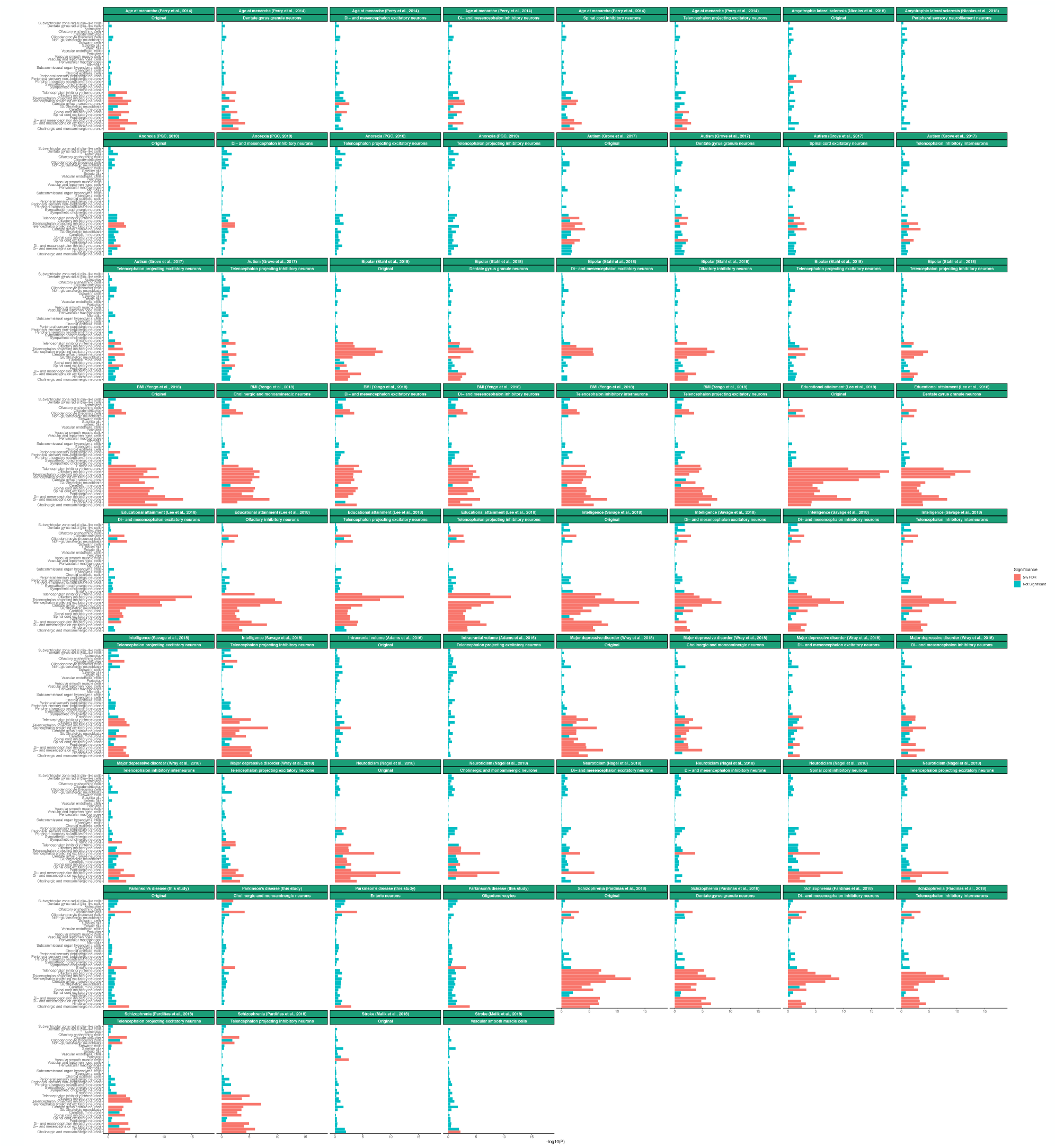
Conditional analysis results for brain related traits. Conditional analysis results using MAGMA are shown for up to the 5 most associated cell types (if at least 5 cell types were significant at a 5% false discovery rate in the original analysis. The color indicates if the cell type is significant at a 5% false discovery rate and the label indicates the cell type the association analysis is being conditioned on.

**Figure S13:**
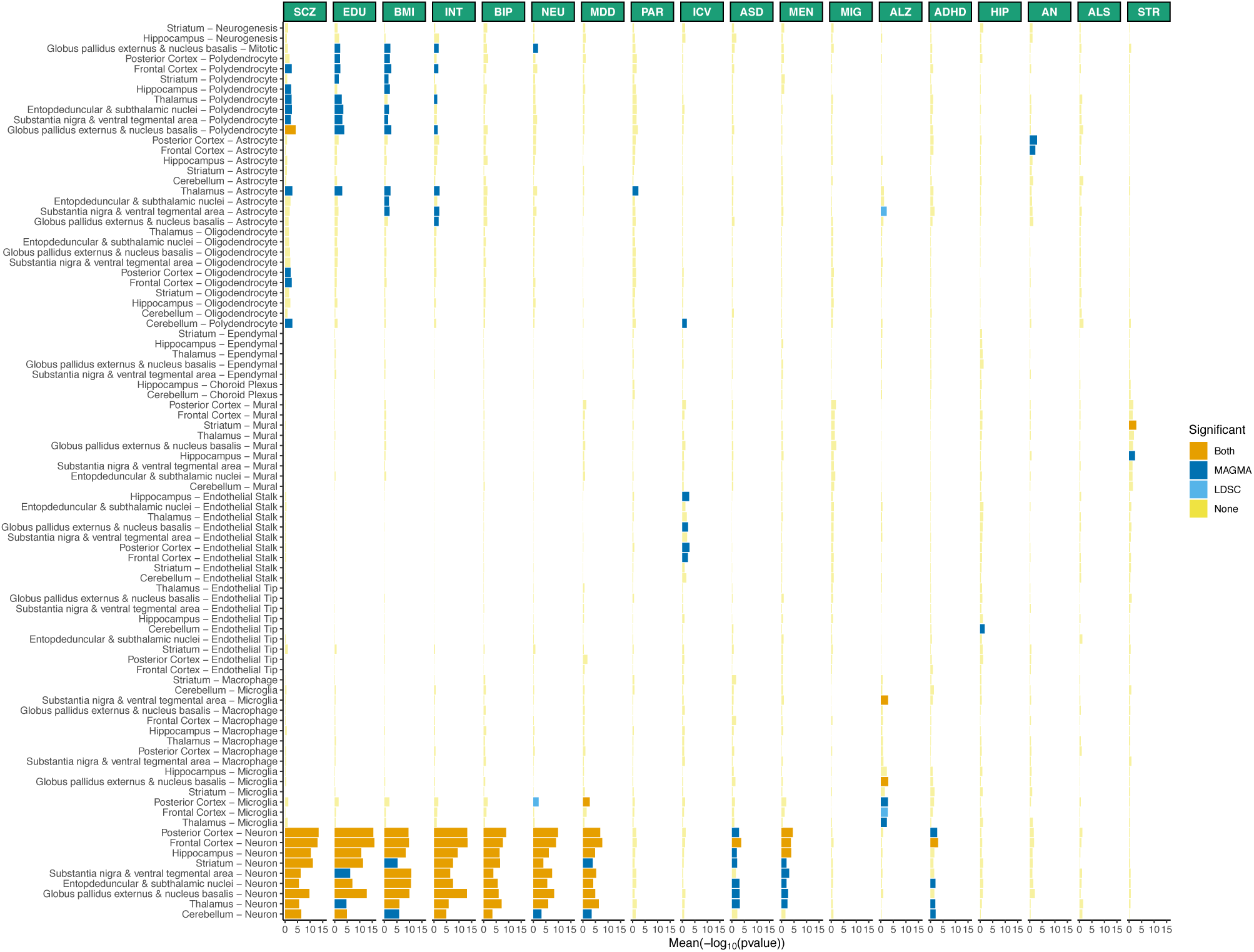
Replication of cell type – trait associations in 88 cell types from 9 different brain regions. The mean strength of association (−log_10_P) of MAGMA and LDSC is shown and the bar color indicates whether the cell type is significantly associated with both methods, one method or none (significance threshold: 5% false discovery rate). SCZ (schizophrenia), EDU (educational attainment), INT (intelligence), BMI (body mass index), BIP (bipolar disorder), NEU (neuroticism), PAR (Parkinson’s disease), MDD (Major depressive disorder), MEN (age at menarche), ICV (intracranial volume), ASD (autism spectrum disorder), STR (stroke), AN (anorexia nervosa), MIG (migraine), ALS (amyotrophic lateral sclerosis), ADHD (attention deficit hyperactivity disorder), ALZ (Alzheimer’s disease), HIP (hippocampal volume).

**Figure S14:**
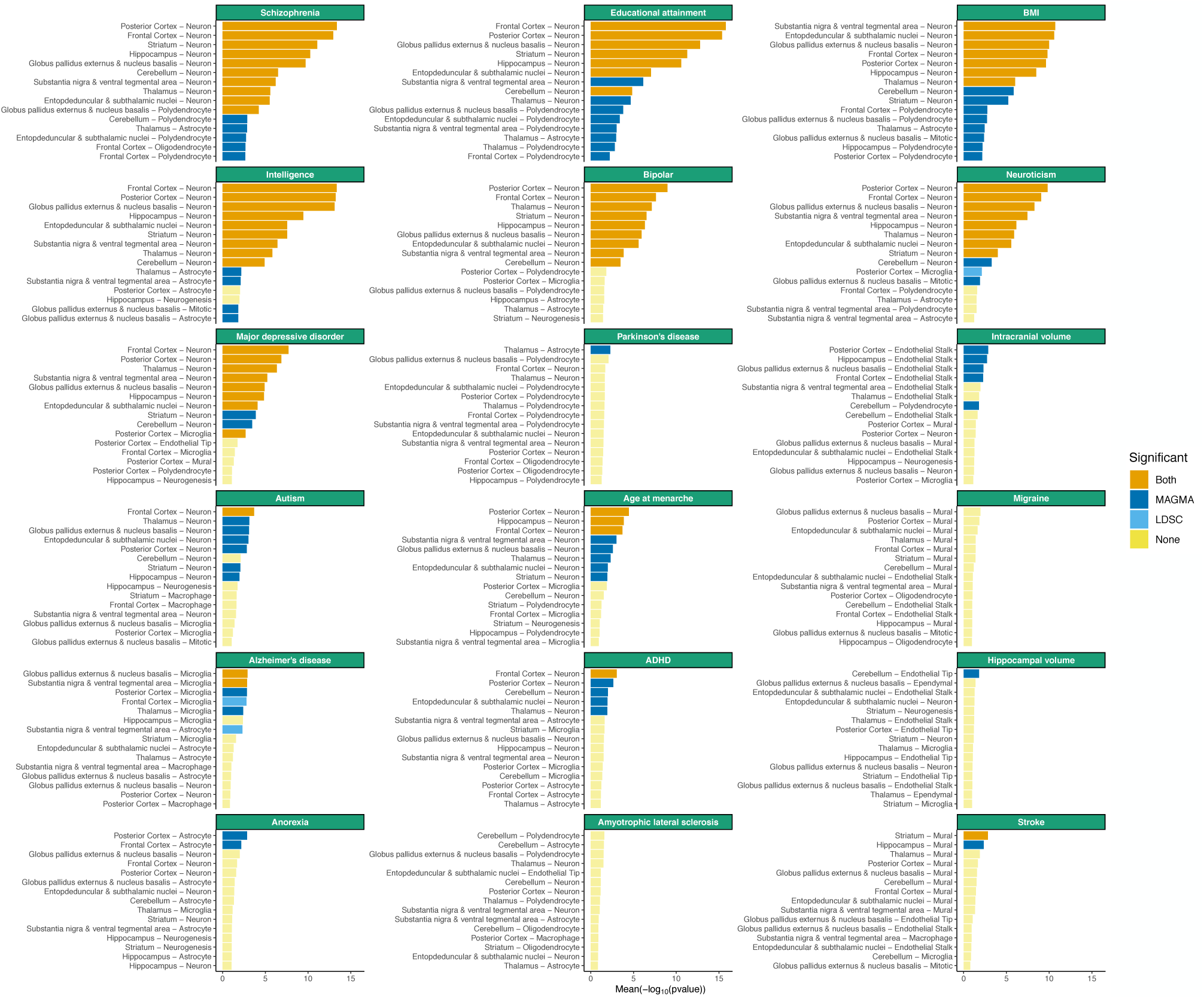
Top associated cell types with brain related traits among 88 cell types from 9 different brain regions. The mean strength of association (−log_10_P) of MAGMA and LDSC is shown for the 15 top cell types for each trait. The bar color indicates whether the cell type is significantly associated with both methods, one method or none (significance threshold: 5% false discovery rate).

**Figure S15:**
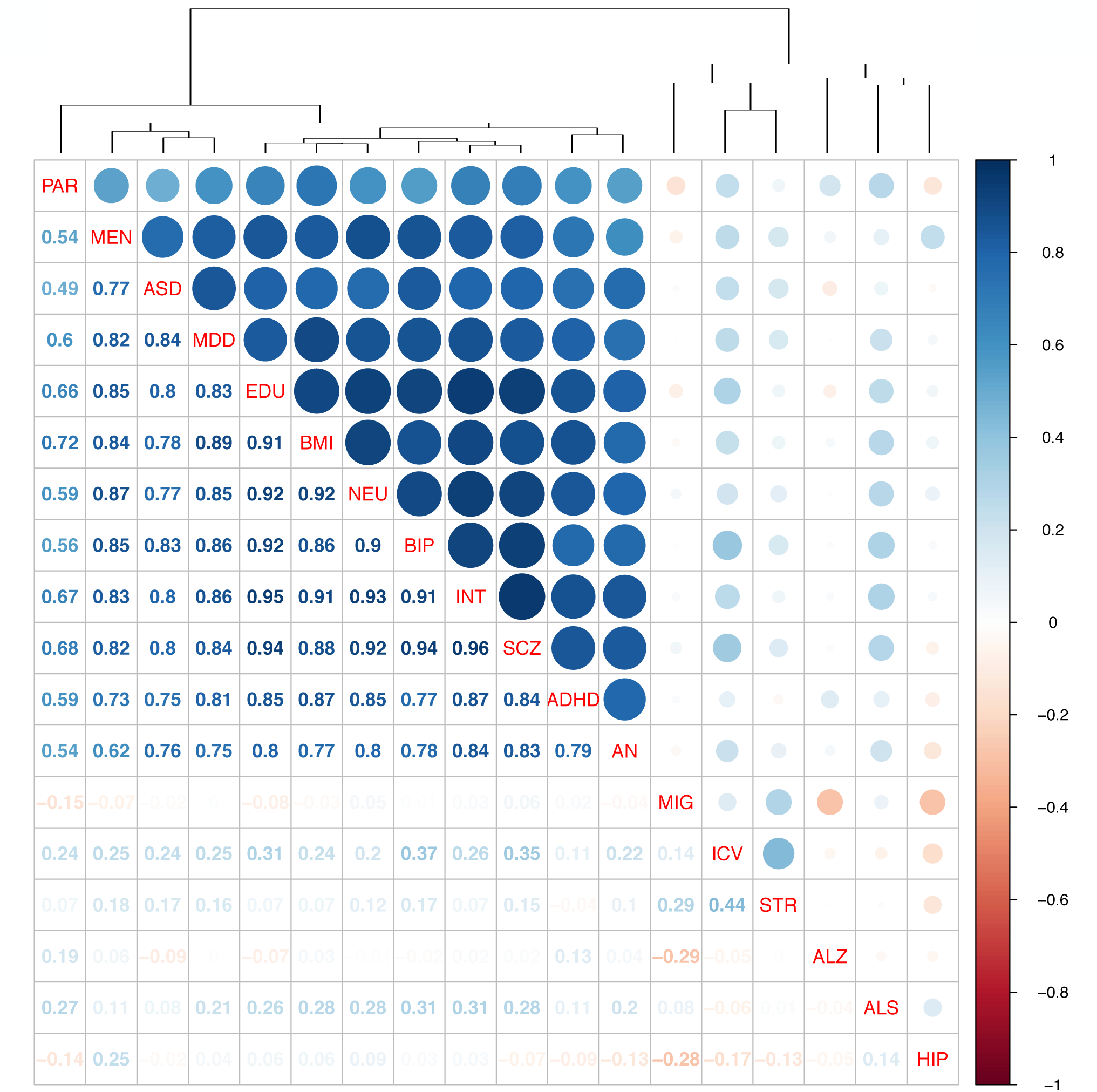
Correlation in cell type associations across traits in a replication data set (88 cell types, 9 brain regions). Spearman rank correlations for cell types associations (−log_10_P) across traits are shown. SCZ (schizophrenia), EDU (educational attainment), INT (intelligence), BMI (body mass index), BIP (bipolar disorder), NEU (neuroticism), PAR (Parkinson’s disease), MDD (Major depressive disorder), MEN (age at menarche), ICV (intracranial volume), ASD (autism spectrum disorder), STR (stroke), AN (anorexia nervosa), MIG (migraine), ALS (amyotrophic lateral sclerosis), ADHD (attention deficit hyperactivity disorder), ALZ (Alzheimer’s disease), HIP (hippocampal volume).

**Figure S16:**
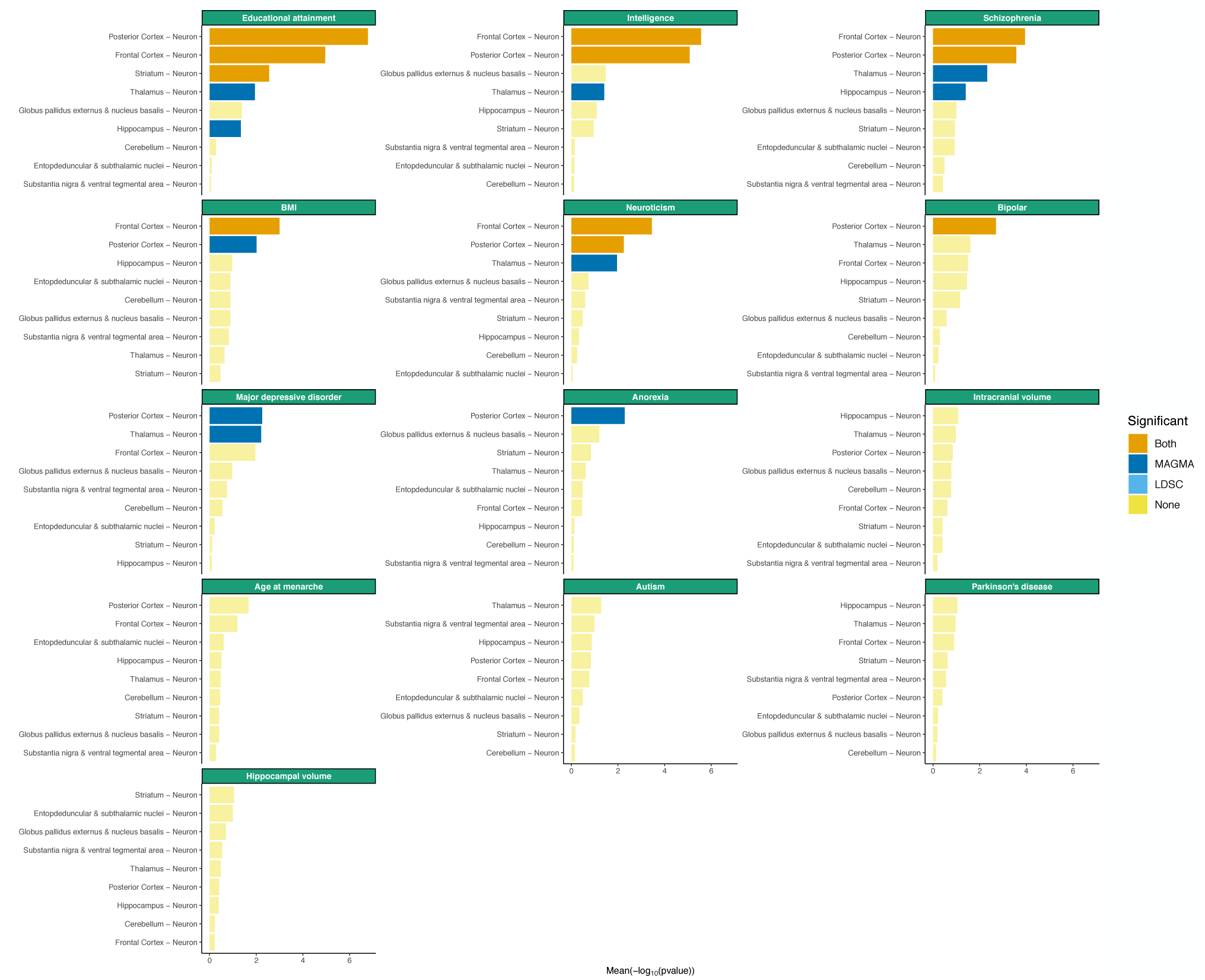
Associations of brain related traits with neurons from 9 different brain regions. Trait – neuron association are shown for neurons of the 9 different brain regions. The specificity metrics were computed only using neurons. The mean strength of association (−log_10_P) of MAGMA and LDSC is shown and the bar color indicates whether the cell type is significantly associated with both methods, one method or none (significance threshold: 5% false discovery rate).

**Figure S17:**
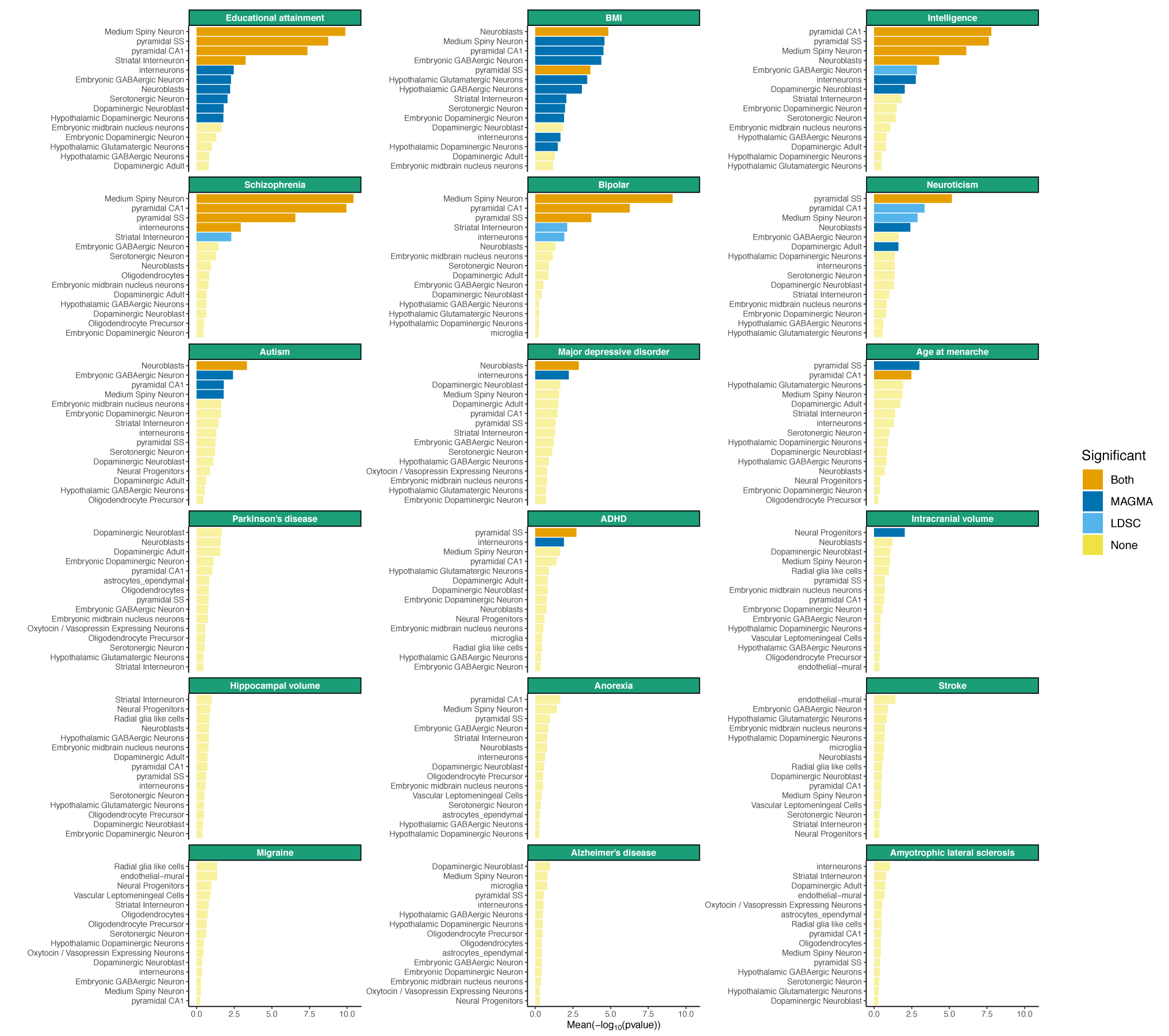
Top associated cell types with brain related traits among 24 cell types from 5 different brain regions. The mean strength of association (−log_10_P) of MAGMA and LDSC is shown for the 15 top cell types for each trait. The bar color indicates whether the cell type is significantly associated with both methods, one method or none (significance threshold: 5% false discovery rate).

**Figure S18:**
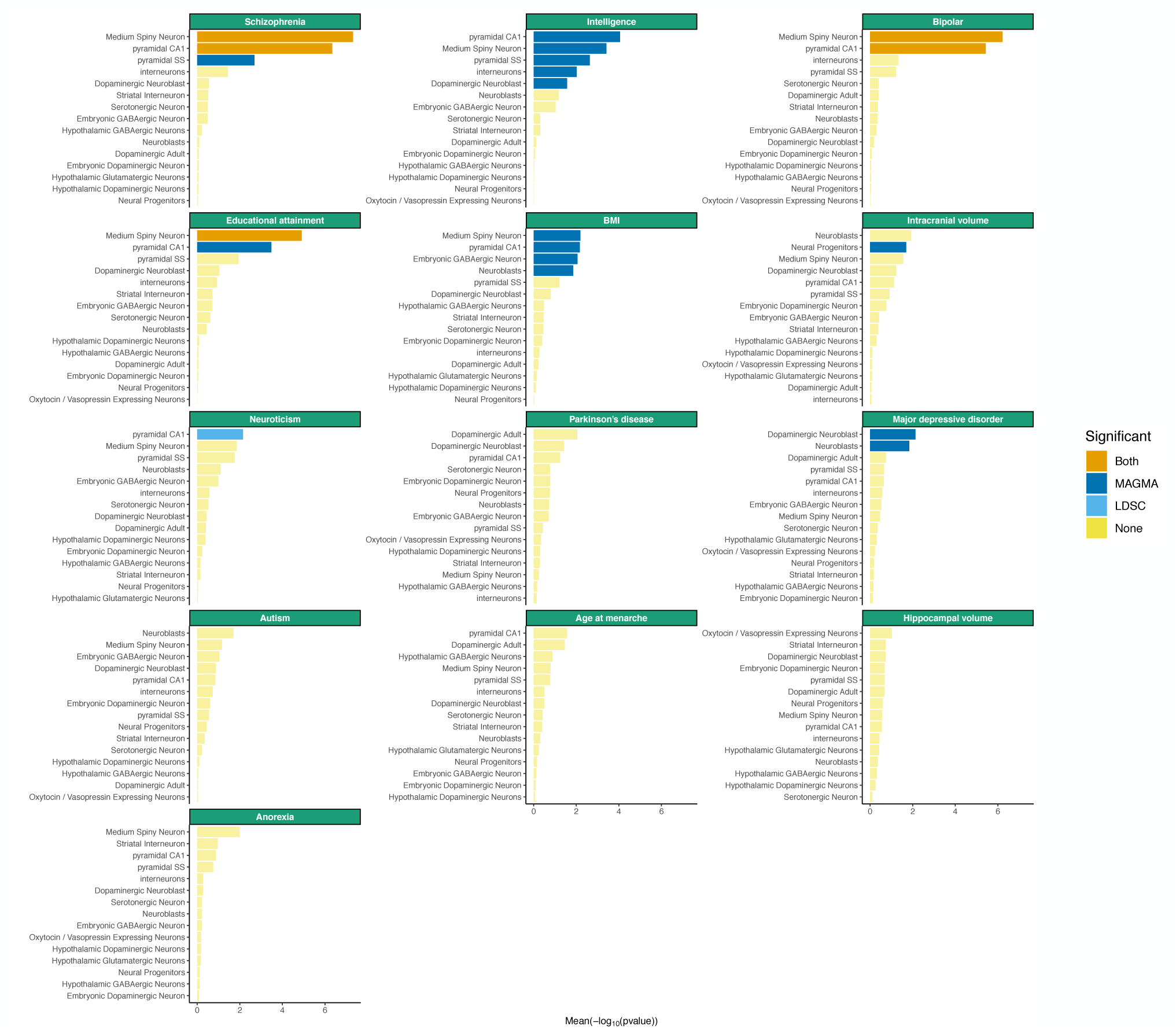
Top associated neurons with brain related traits among 16 neurons from 5 different brain regions. The specificity metrics were computed only using neurons. The mean strength of association (−log_10_P) of MAGMA and LDSC is shown for the top 15 cell types for each trait. The bar color indicates whether the cell type is significantly associated with both methods, one method or none (significance threshold= 5% false discovery rate).

**Figure S19:**
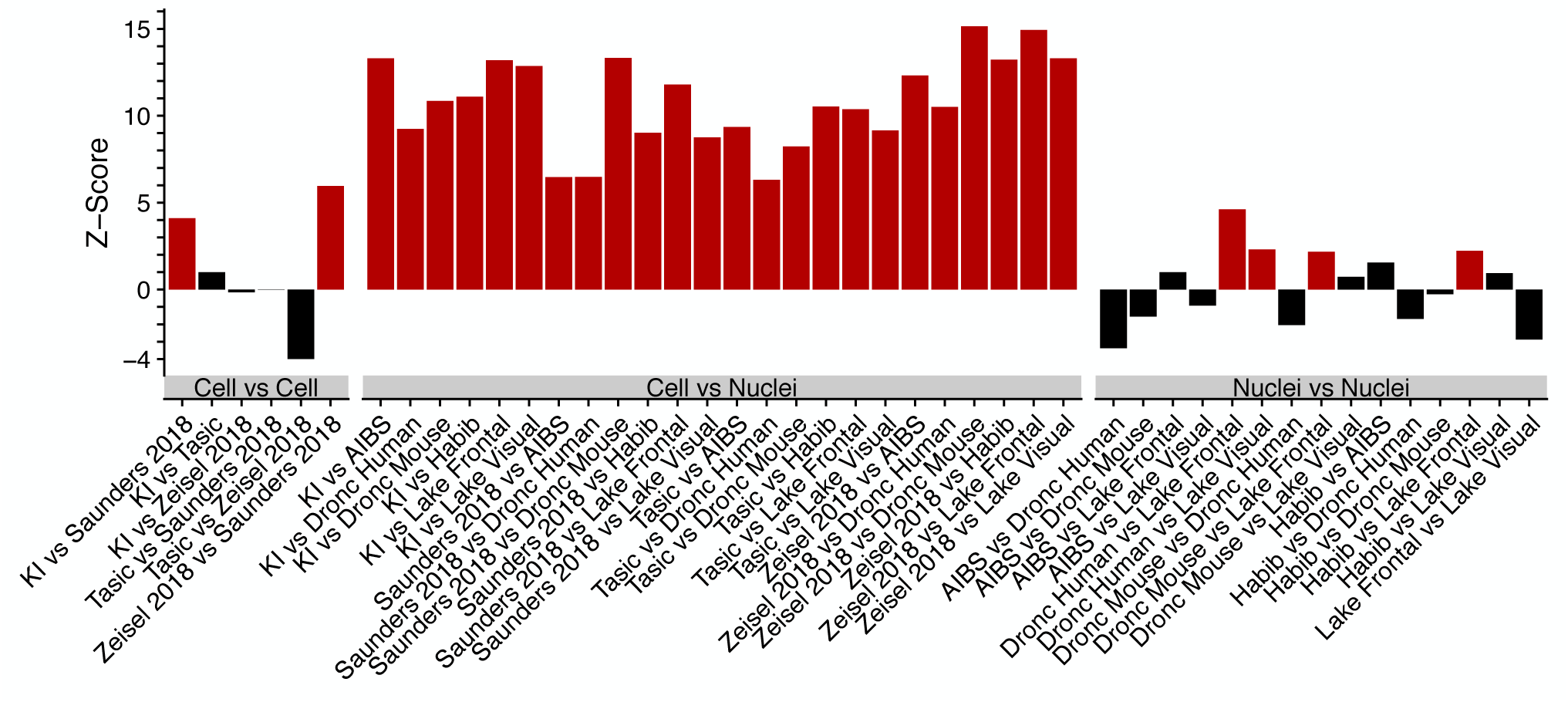
Single nuclei datasets are systematically depleted of dendritically enriched transcripts relative to single-cell datasets. Each bar represents a comparison between two datasets (X versus Y), with the bootstrapped z-scores representing the extent to which dendritically enriched transcripts^96^ have lower specificity for pyramidal neurons in dataset Y relative to that in dataset X. Larger z-scores indicate greater depletion of dendritically enriched transcripts, and red bars indicate a statistically significant depletion (P < 0.05, by bootstrapping).

**Figure S20:**
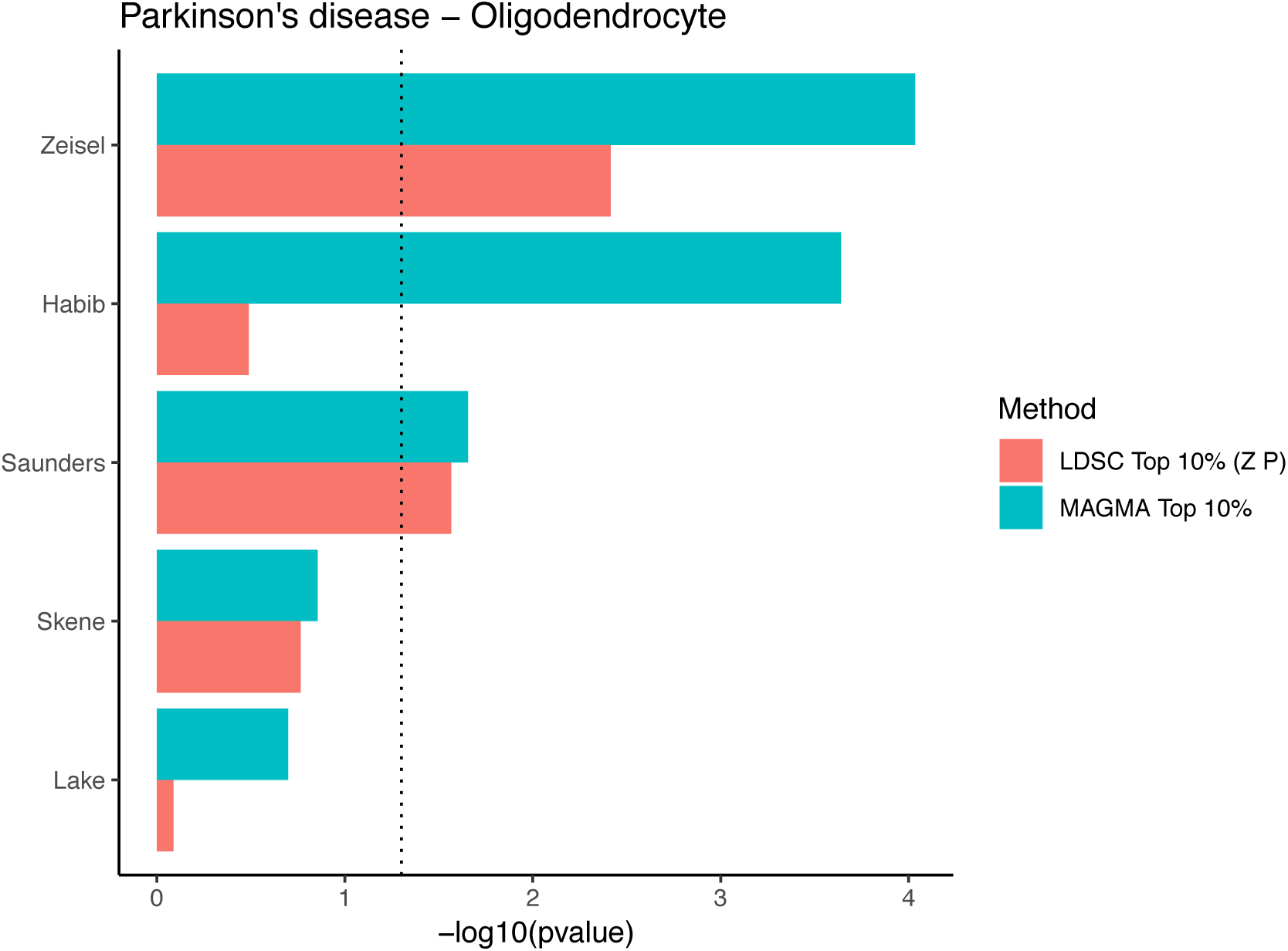
Association of Parkinson’s disease with oligodendrocytes in the different datasets. The dotted line indicated the nominal significance threshold (P=0.05)

**Figure S21:**
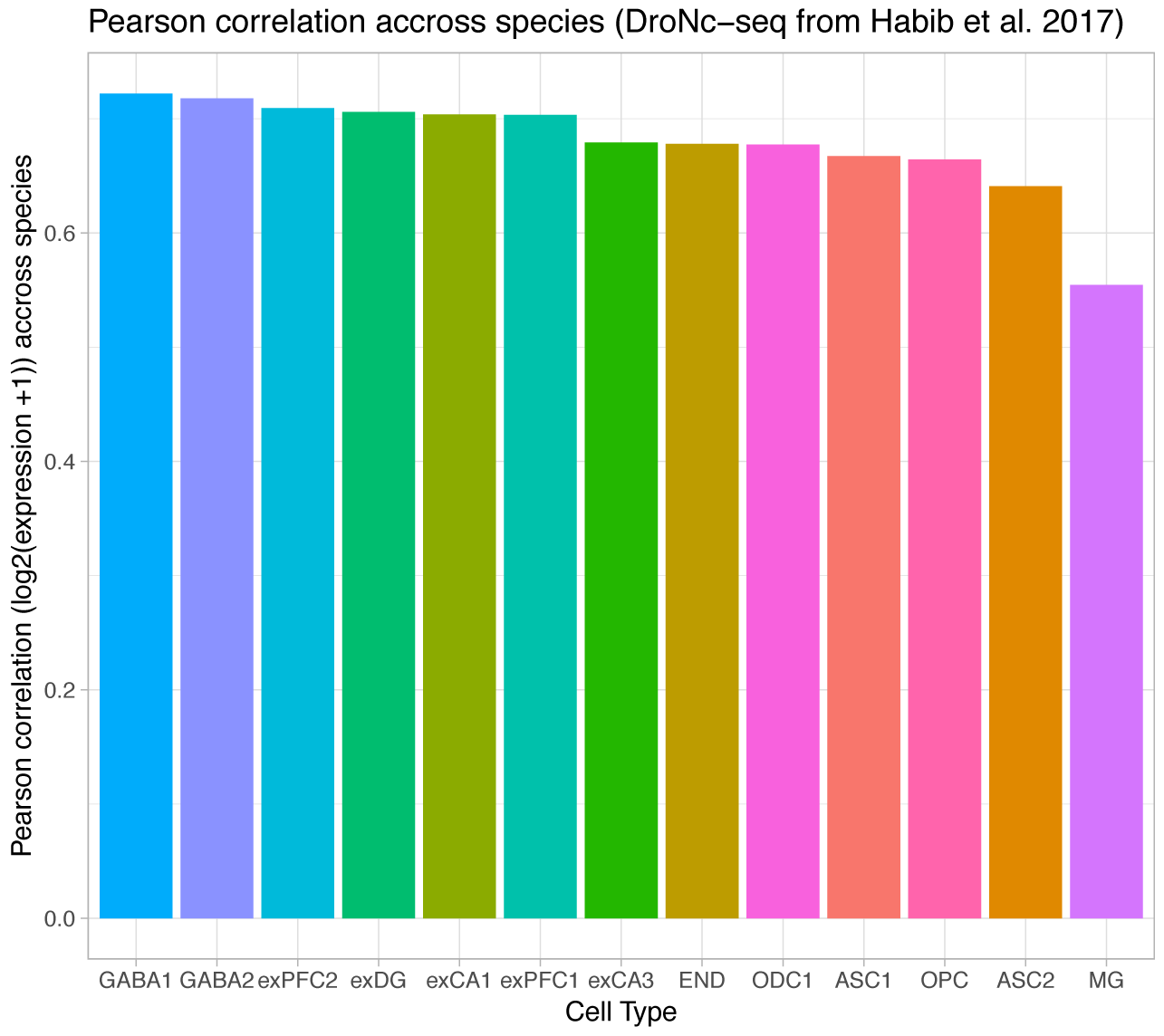
Gene expression correlation within cell type across species. Pearson correlation of gene expression (log_2_(expression) +1) between mouse and human cell types with matching names (from Habib et al. 2017 ^42^).

**Figure S22:**
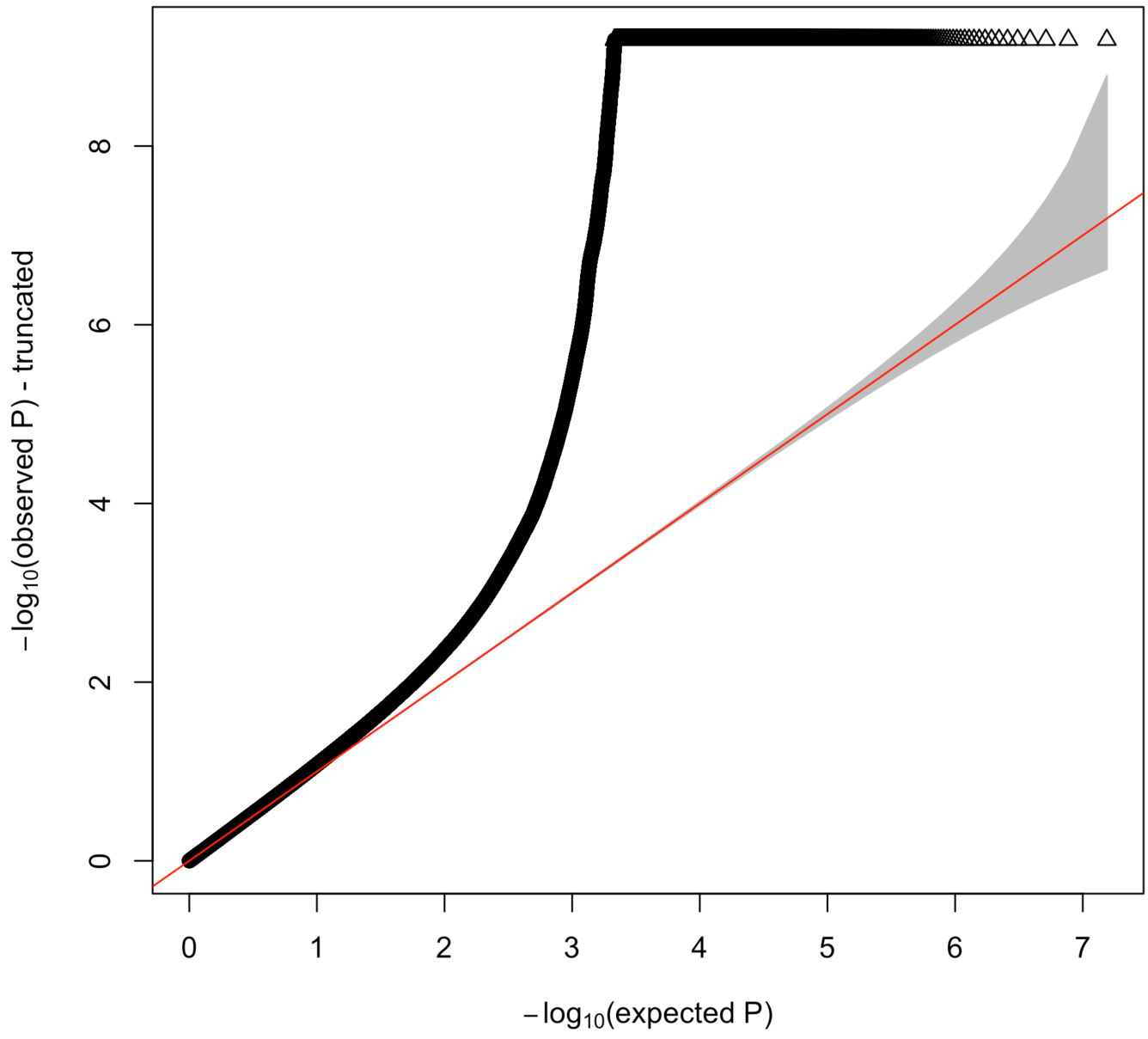
Quantile-quantile plot of Parkinson’s disease meta-analysis. Quantile-quantile plot of the meta-analyzed pvalues for Parkinson’s disease. The y-axis is truncated for clarity. The grey zone around the red line represents the 95% confidence interval for the null distribution.

**Figure S23:**
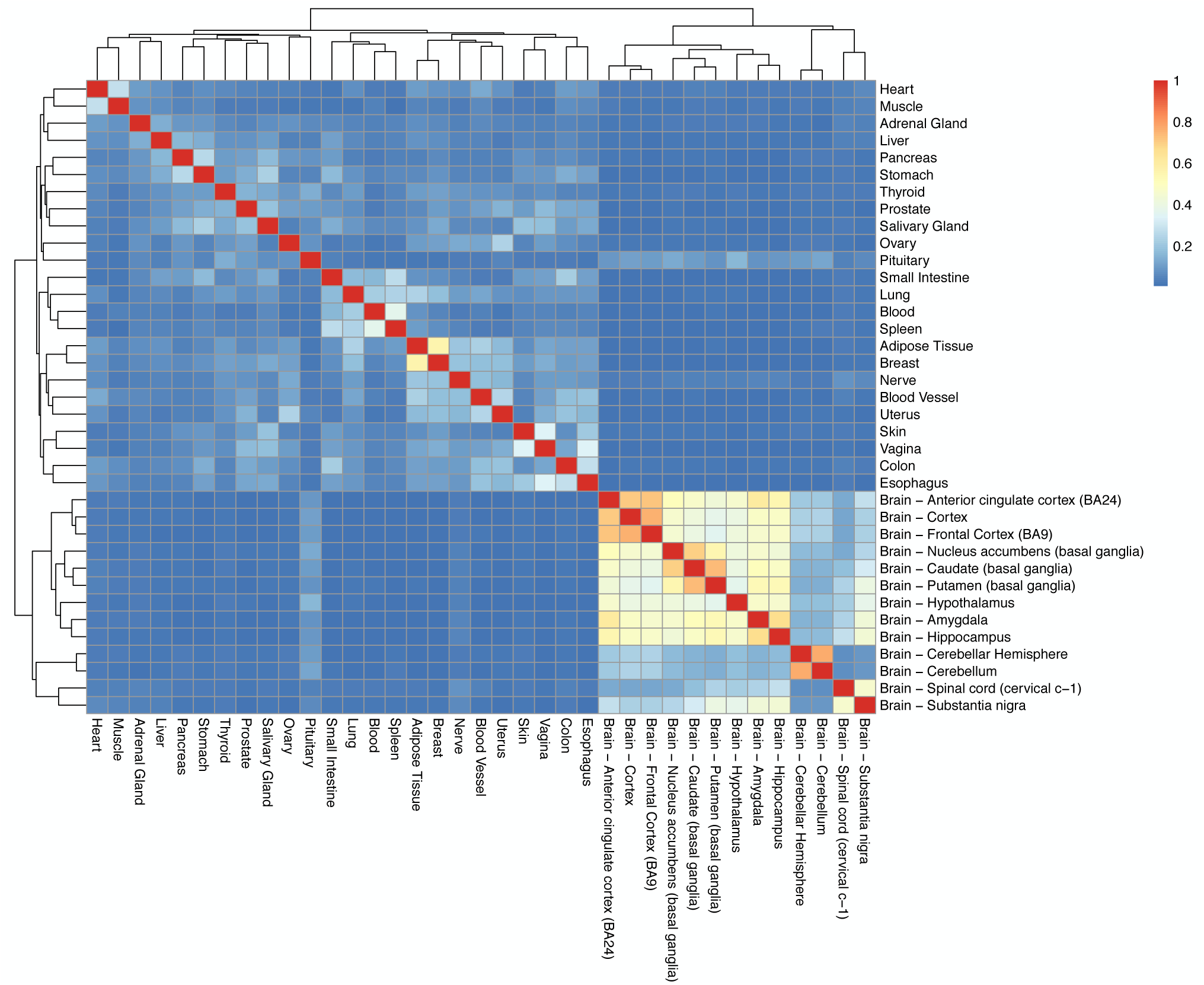
Jaccard index for the top 10% most specific genes in each tissue in the GTEx dataset. Jaccard index were calculated between the top 10% most specific genes in each tissue from the GTEx dataset.

**Figure S24:**
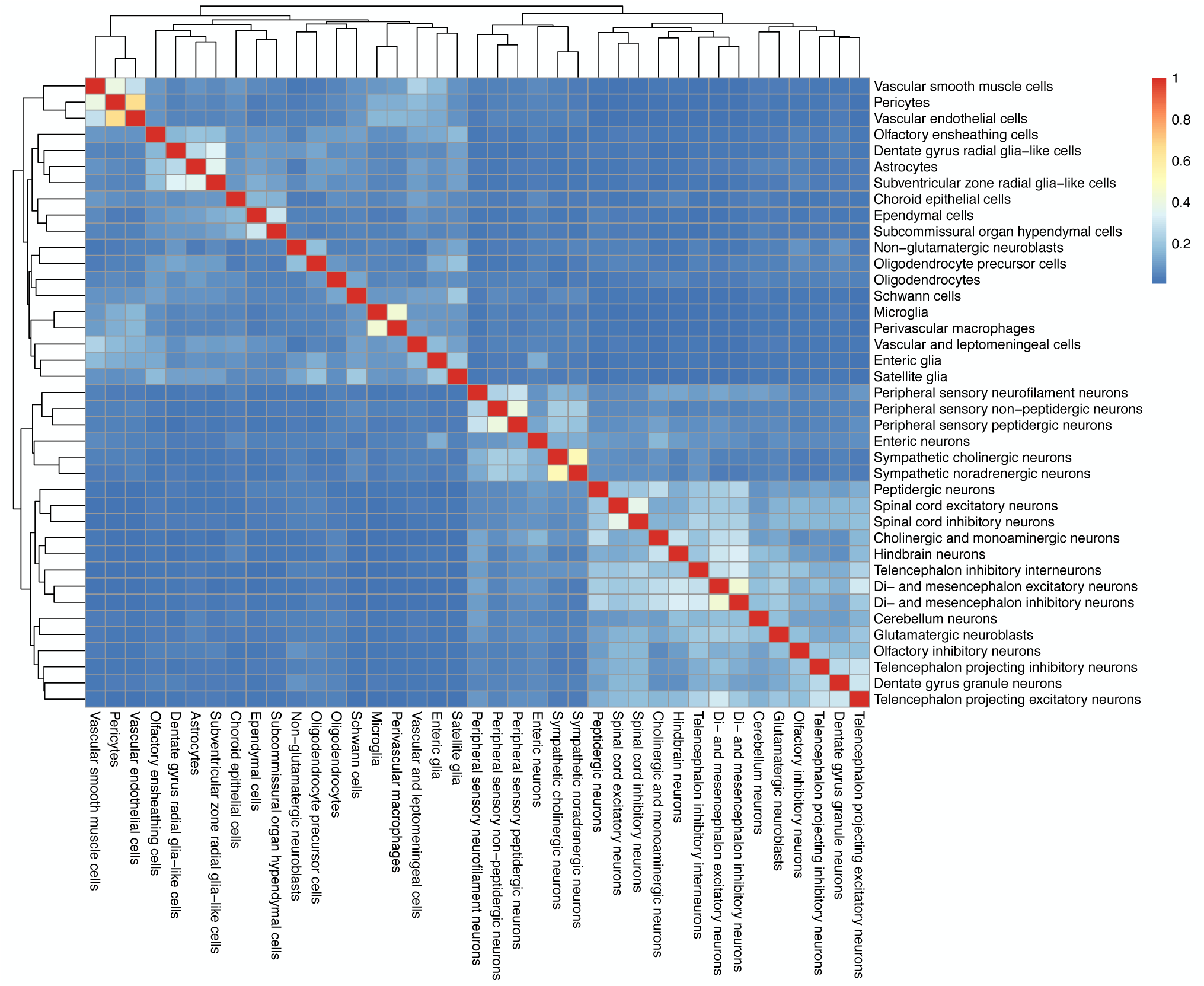
Jaccard index for the top 10% most specific genes in each cell type in the mouse nervous system (Zeisel et al. 2018). Jaccard index were calculated between the top 10% most specific genes in each cell type from the mouse nervous system (Zeisel et al. 2018).

**Figure S25:**
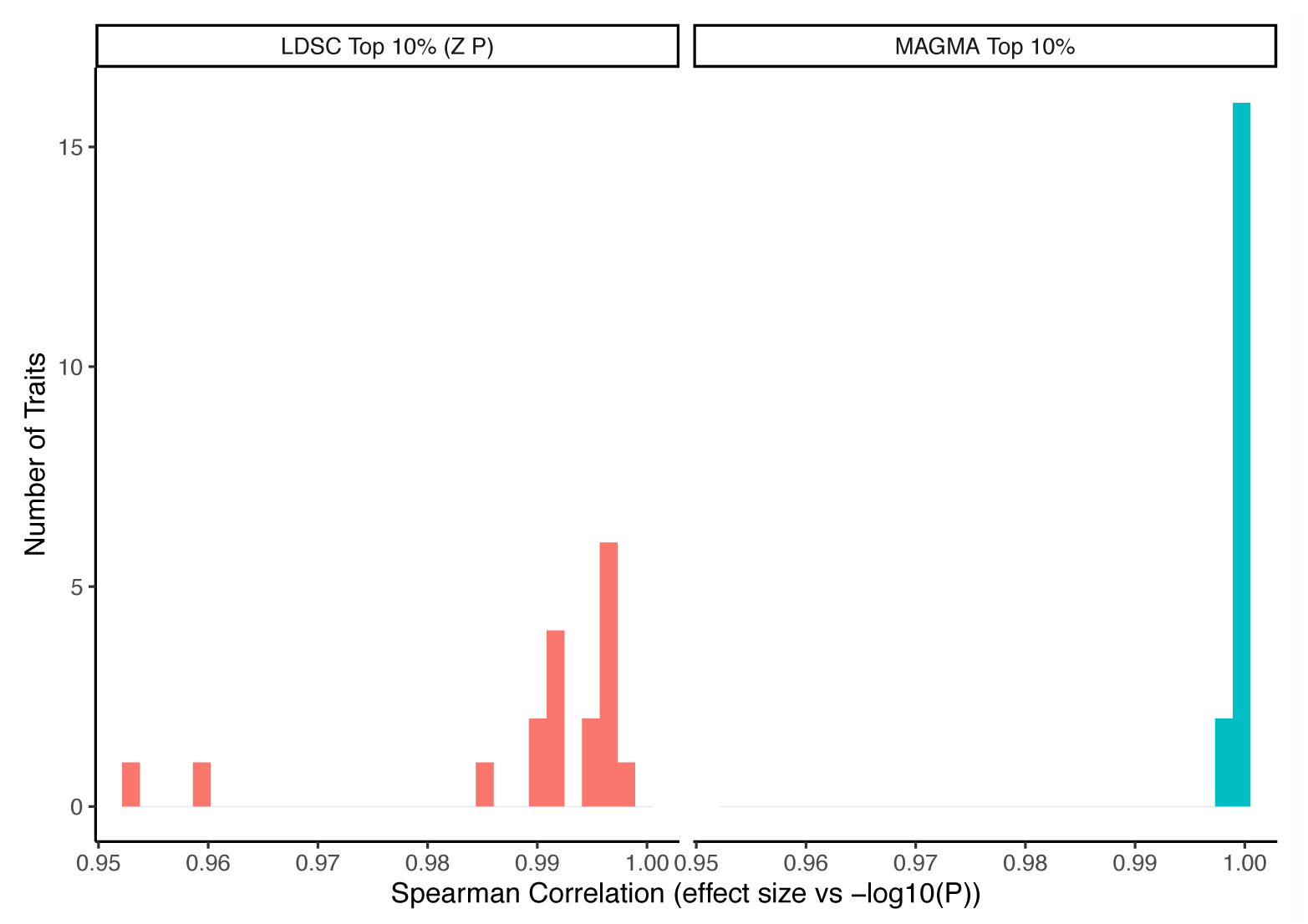
Correlation between beta coefficient and significance level. Histograms of the spearman rank correlations between effect size (beta coefficient) and significance (−log_10_P) computed for each trait in the Zeisel dataset. The effect sizes are strongly correlated with the significance level of the cell type with values ranging from 0.999 to 1 using MAGMA and 0.953 to 1 with LDSC.

**Figure S26:**
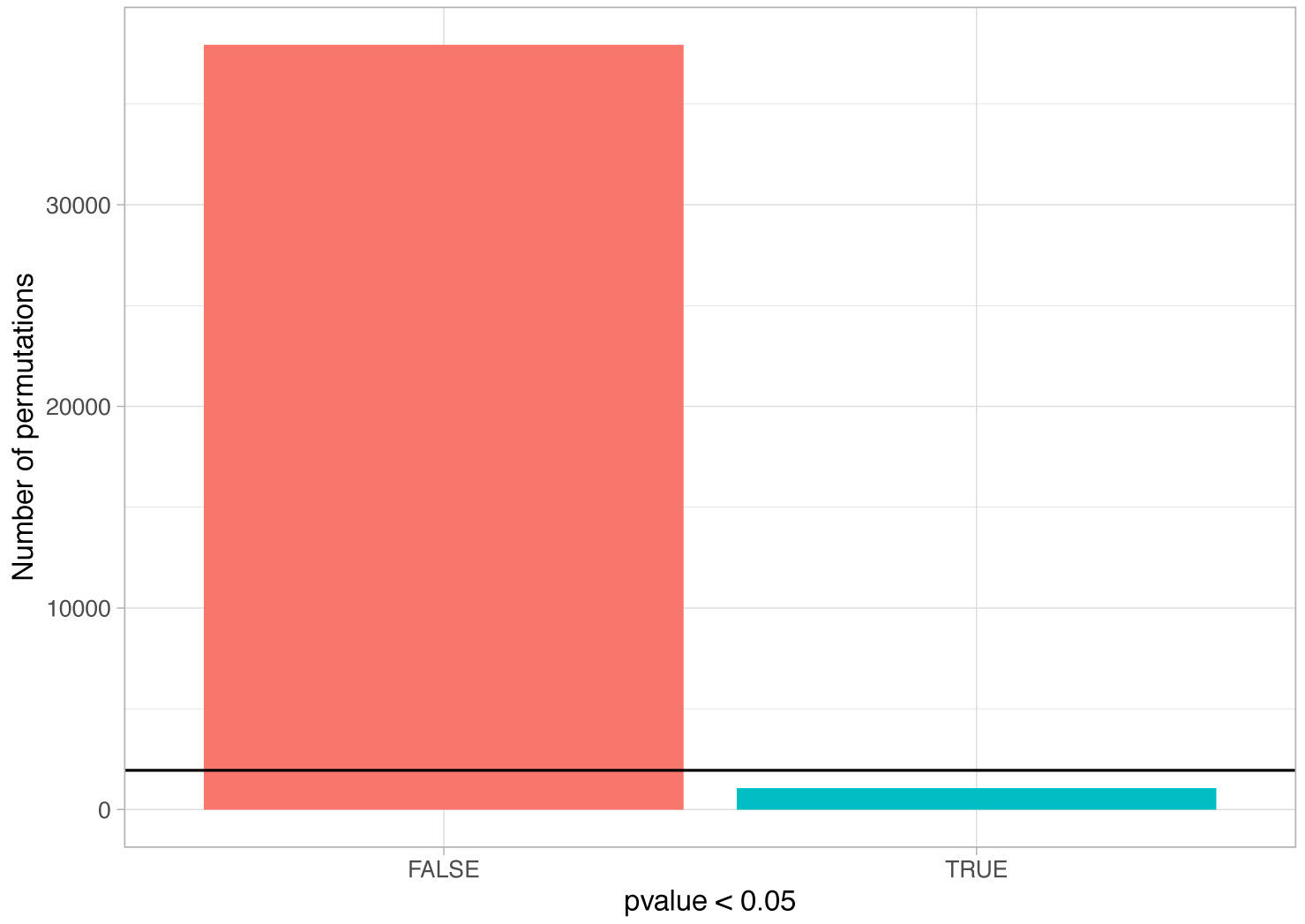
Number of MAGMA associations with P<0.05 using permuted gene-level genetic associations. Gene labels were randomly permuted a thousand times for the schizophrenia MAGMA gene-level genetic associations (39 cell types * 1000 permuted labels=39,000 associations with permuted gene labels). The number of permutations with P < 0.05 is shown in blue. The black horizontal bar shows expected number of random associations with P < 0.05 (39,000*0.05=1950).

**Figure S27:**
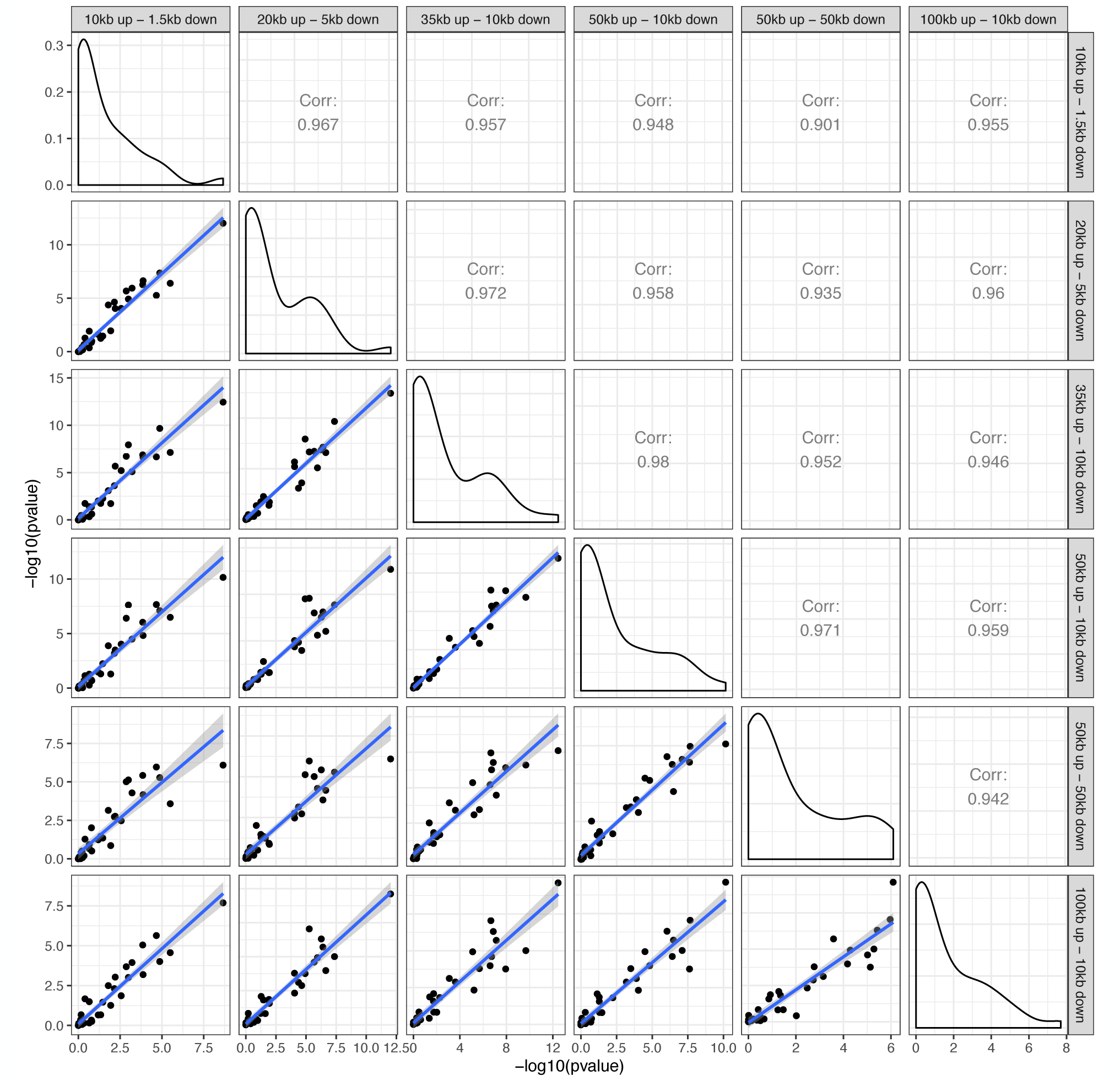
Correlation in schizophrenia cell type association strengths with different window sizes using MAGMA. Pearson correlations of the cell type association strength (−log_10_P) across different window sizes using MAGMA. The diagonal shows the distribution of the (−log_10_P) for each window size.

**Figure S28:**
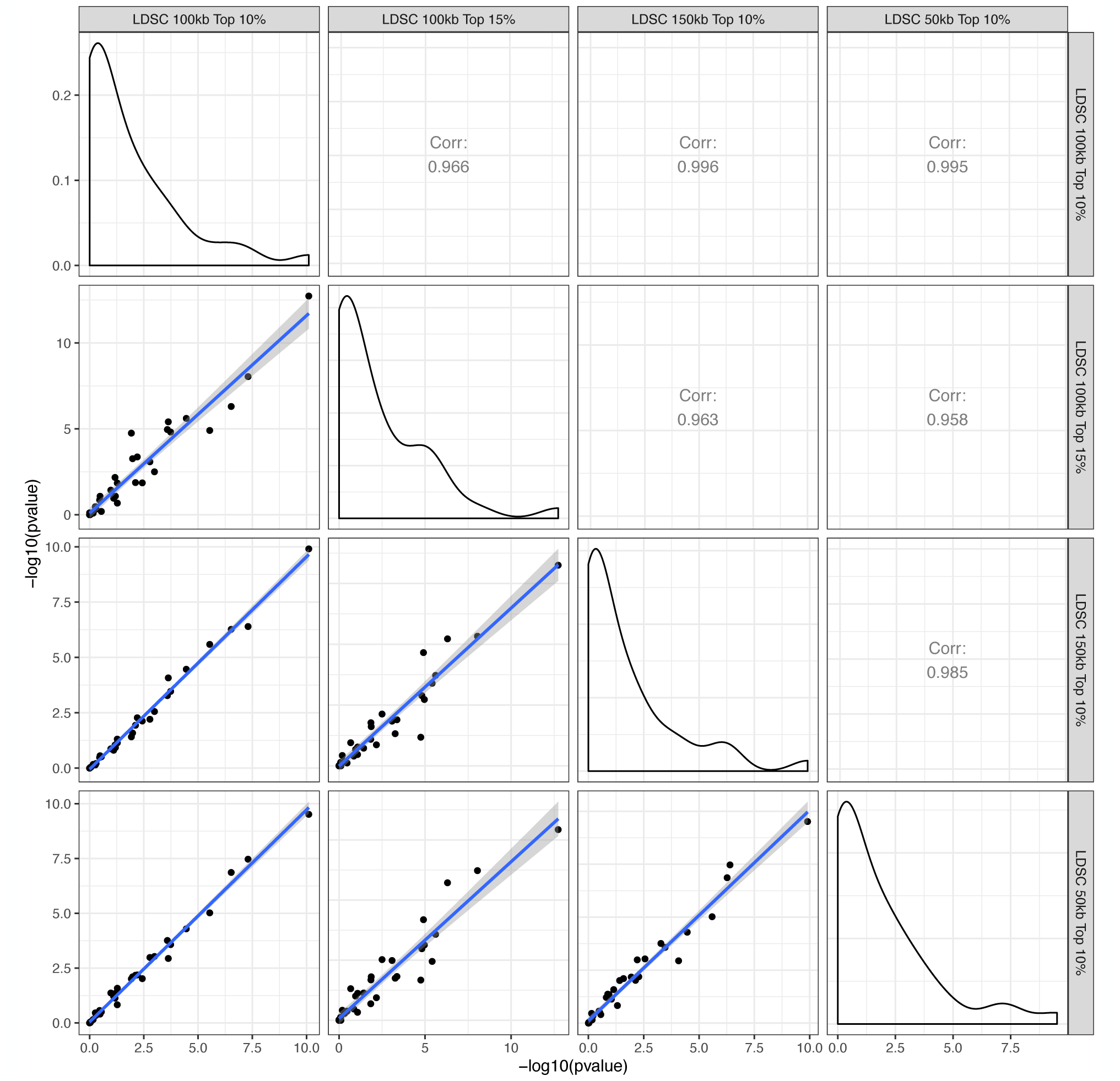
Correlation in schizophrenia cell type association strengths with different window sizes and percentages of most specific genes using LDSC. Pearson correlations of the cell type association strength (−log_10_P) across different window sizes and percentages of most specific genes using LDSC. The diagonal shows the distribution of the (−log_10_P) for the cell type associations using different parameters.

**Figure S29:**
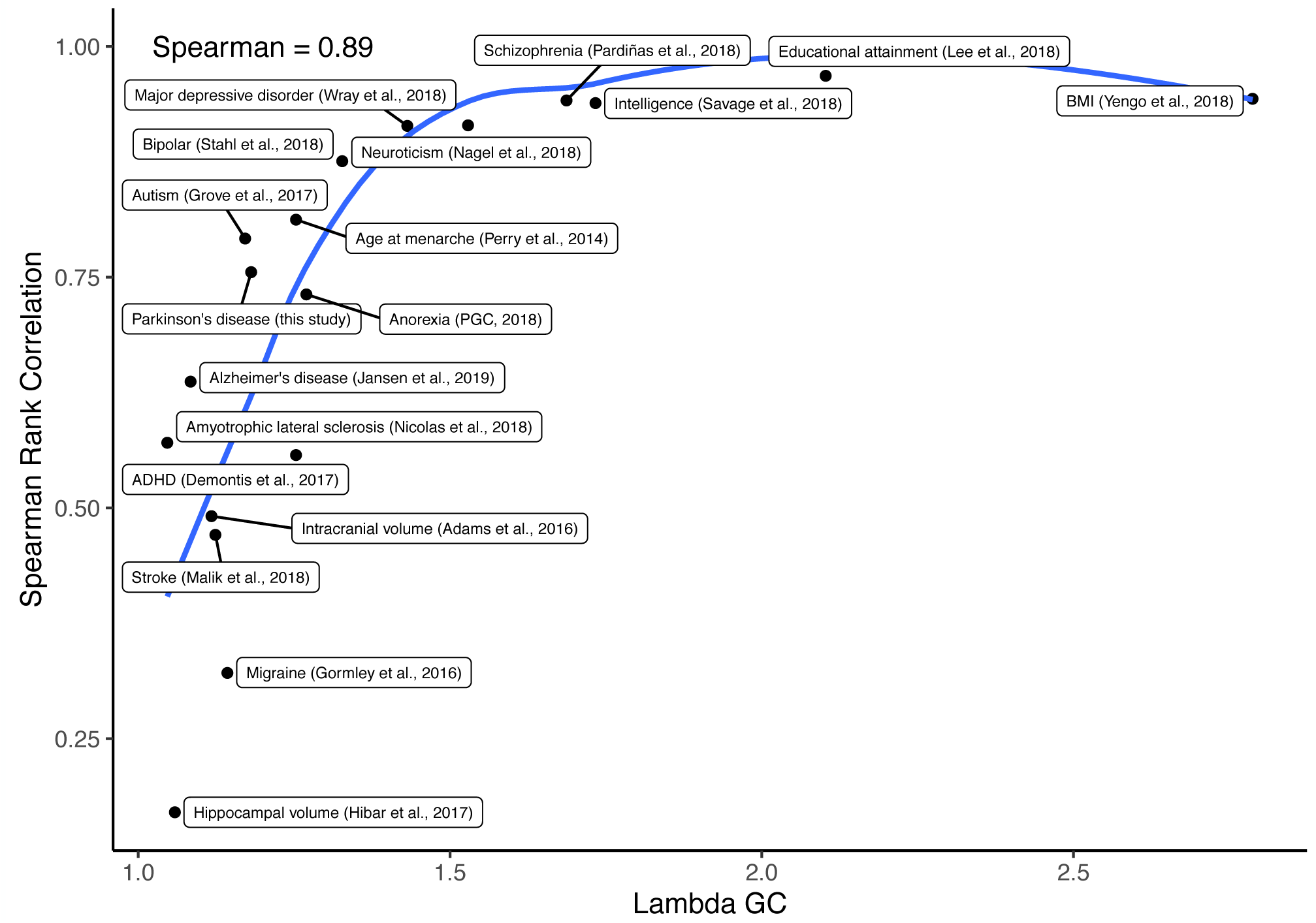
Correlation between *λ_GC_* and similarity in cell type ordering between MAGMA and LDSC. LDSC^101^ was used to obtain *λ_GC_* (a measure of the deviation of the GWAS statistics from the expected) for each GWAS. Spearman rank correlation was used to test for similarity in association strength (− log_10_P) between MAGMA and LDSC for each GWAS among 39 cell types from the nervous system.

**Figure S30:**
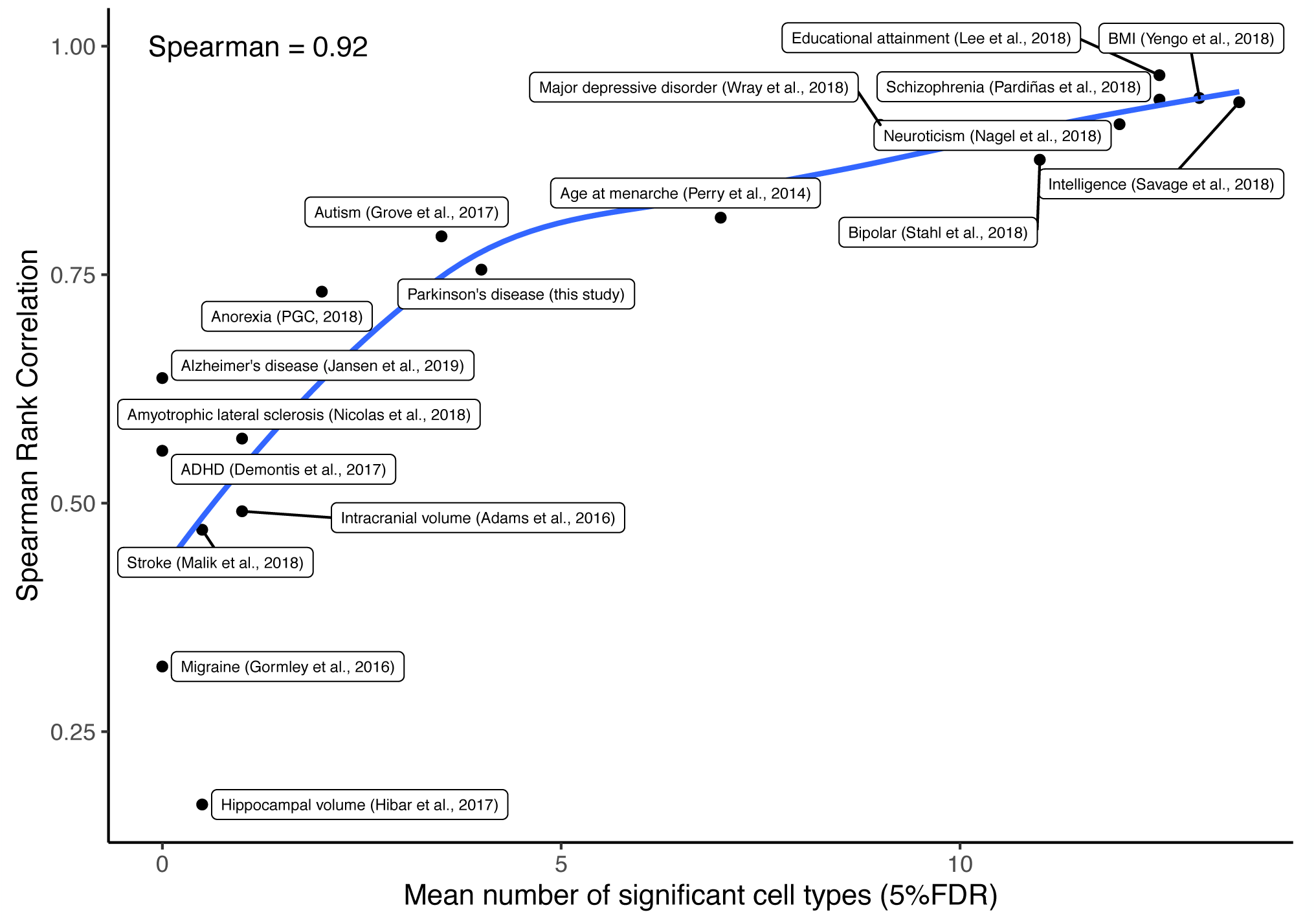
Correlation between mean number of significant cell types and similarity in cell type ordering between MAGMA and LDSC. The mean number of cell types was obtained by taking the average of the number of cell types that were significantly associated with each trait (FDR<5%) using MAGMA and LDSC. Spearman rank correlation was used to test for similarity in association strength (−log_10_P) between MAGMA and LDSC among 39 cell types from the nervous system.

**Figure S31:**
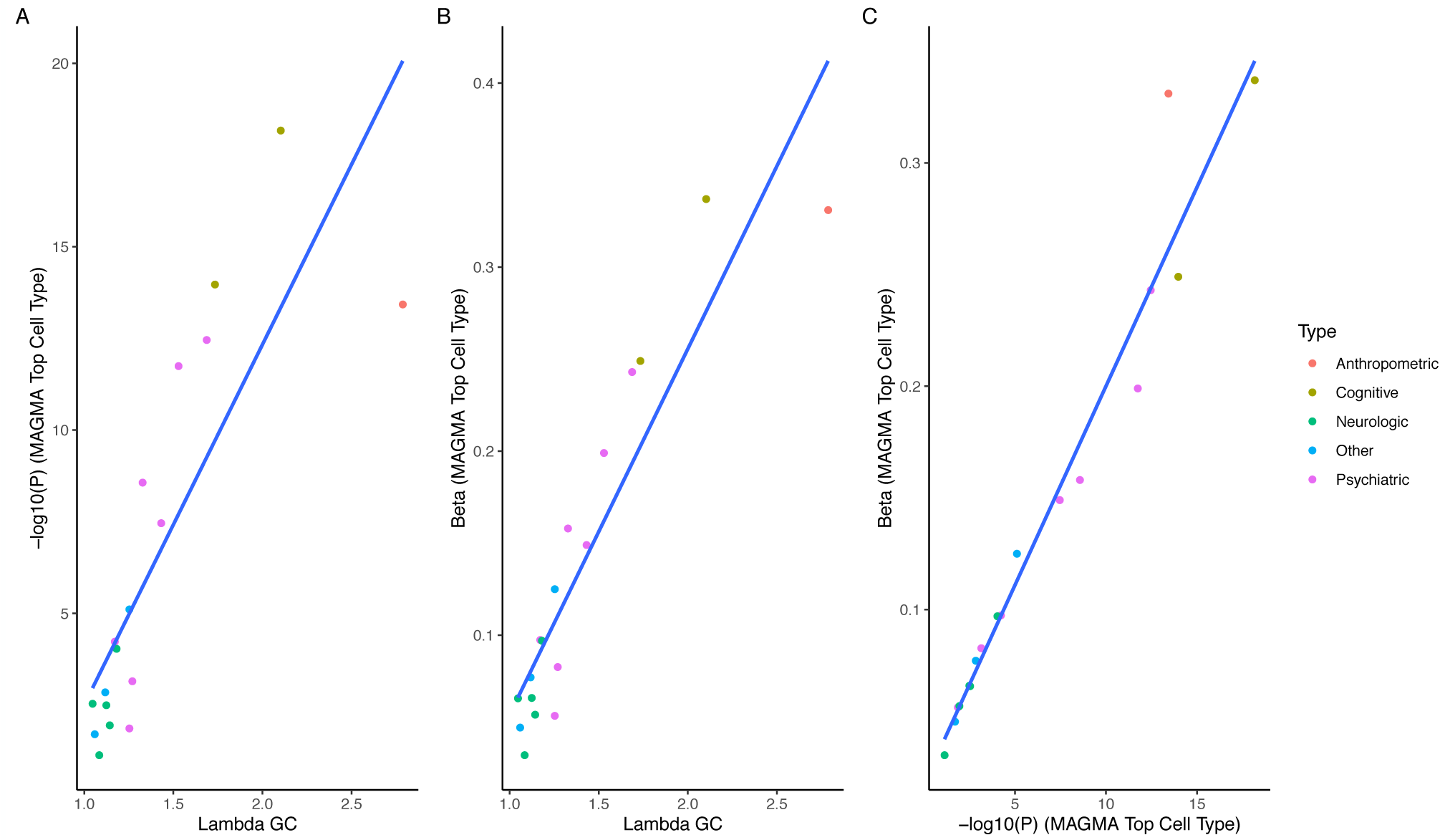
The GWAS λ_GC_ is correlated with the strength of association of the top cell type in the Zeisel dataset. Scatter plot of the λ _GC_ (median of chi-squared test statistics divided by expected median of the chi-squared distribution) of each GWAS vs the strength of association of the top Zeisel cell type associated with the trait (−log_10_(P_MAGMA_)). Spearman correlation=0.88 (**A**). Scatter plot of the λ_GC_ of each GWAS vs the effect size of the top Zeisel cell type associated with the trait (− log_10_(P_MAGMA_)). Spearman correlation=0.9 (**B**). Scatter plot of the strength of association of the top Zeisel cell type (−log_10_(P_MAGMA_)) of each GWAS vs the effect size of the top Zeisel cell type. Spearman correlation=0.996 (**C**).

